# Changes in interactions over ecological time scales influence single cell growth dynamics in a metabolically coupled marine microbial community

**DOI:** 10.1101/2022.02.08.479118

**Authors:** Michael Daniels, Simon van Vliet, Martin Ackermann

**Affiliations:** Department of Environmental Systems Sciences, Microbial Systems Ecology Group, Institute of Biogeochemistry and Pollutant Dynamics, ETH-Zurich, Zurich, Switzerland; Department of Environmental Microbiology, Eawag: Swiss Federal Institute of Aquatic Sciences, Duebendorf, Switzerland; Interdisciplinary PhD Program Systems Biology, ETH-Zurich & University of Zurich, Zurich, Switzerland; Biozentrum, University of Basel, Switzerland

## Abstract

Microbial communities thrive in almost all habitats on earth. Within these communities, cells interact through the release and uptake of metabolites. These interactions can have synergistic or antagonistic effects on individual community members. The collective metabolic activity of microbial communities leads to changes in their local environment. As the environment changes over time, the nature of the interactions between cells can change. We currently lack understanding of how such dynamic feedbacks affect the growth dynamics of individual microbes and of the community as a whole. Here we study how interactions mediated by the exchange of metabolites through the environment change over time within a simple marine microbial community. We used a microfluidic-based approach that allows us to disentangle the effect cells have on their environment from how they respond to their environment. We found that the interactions between two species - a degrader of chitin and a cross-feeder that consumes metabolic by-products - changes dynamically over time as cells modify their environment. Cells initially interact positively and then start to compete at later stages of growth. Our results demonstrate that interactions between microbes are not static and depend on the state of the environment, emphasizing the importance of disentangling how modifications of the environment affects species interactions. This experimental approach can shed new light on how interspecies interactions scale up to community level processes in natural environments.

## Introduction

Microbes play an important role in virtually all natural environments, from the soil to the human host. They contribute to the global cycling of the elements (1), our health (2), as well as a plethora of industrial processes (3). We are beginning to understand the importance of species diversity and interspecies interactions within microbial communities for ecosystem processes (4). Yet, we currently lack quantitative insight into the exact mechanisms of how interspecies interactions modulate the growth of individual community members, and how the growth of individual community members scales up to determine the activity and growth of the community. Such insight would be needed to understand the role microbial communities play in global scale processes.

Species in microbial consortia often perform metabolic functions collectively, in the sense that catabolic and anabolic processes are distributed across different species (5). These species interact with each other through the exchange of metabolites to form complete metabolic pathways together. The distribution of catabolic pathways across species is often observed for complex substrates whose degradation involves several enzymatic reactions (6). In natural environments a major resource for microbial communities are natural polymers (7). These polymers, e.g. polysaccharides, are integral structural components of multicellular organisms, such as cellulose in plants and chitin in the exoskeletons of arthropods. These components are released into the environment upon the death of these organisms. A complex community of interacting species is typically formed on these polymers (8). Specialized bacteria cleave polymers via extracellular enzymes into accessible subunits, i.e. sugar mono- or oligomers that are small enough to be imported into the cell (9). This process builds the foundation of a trophic cascade that leads to the eventual remineralization of natural polymers. Some bacterial cells without the ability to degrade polymers are able to take up the degradation products from the surrounding environment (10). These so-called consumers thus directly benefit from the presence of degraders. Other microbial species that lack the necessary enzymes for polymer degradation are additionally unable to catabolize the cleaved degradation products; they rely on excreted metabolic by-products as growth substrates (11). This process is called cross-feeding.

Cross-feeding, or syntrophy, is an interaction in which one organism uses another organism’s by-products as a resource (12) (13). Cross-feeding is widely believed to underlie many mutualistic relationships between microbes, providing a driving force for the maintenance of diversity in microbial communities (14) (15) (16). Cross-feeding can also be beneficial for the species that excretes the by-products. Firstly, removing the product of biochemical reactions increases the reaction rate (17). Here, according to “Le Chatelier’s principle “, the cross-feeding bacteria serves as a sink for the reaction product and drives the reaction forward. Secondly, cross-feeding can allow for the removal of toxic metabolic by-products that inhibit the growth of their producers. The accumulation of toxic waste products in the environment leads to growth impairment (18) (19). This process can be counteracted by their consumption through cross-feeding species in microbial communities. Therefore, it is widely believed that cross-feeding can be beneficial to both species (20).

Interactions between species are generally classified as positive, negative, or neutral and are usually quantified by measuring the difference in yield or growth rate in co-vs monocultures (21). This approach assumes that interactions within communities remain constant during the time period over which they are measured. At ecological scales, microbes transition through various growth phases i.e. lag, exponential, and stationary. By consuming and releasing nutrients, cells dynamically change their environment. In turn, individuals alter their metabolic behavior to adjust to the changes they cause in their surroundings (22). This process can cause metabolic interactions between species to change with time.

Cells respond to changes in the environment that are caused both by their own activity and by the activity of all other community members. This leads to a complex feedback loop where the state of the environment, the metabolic activity of the community members, and the interactions between all these members become interdependent (12). To gain a mechanistic understanding of these complex community dynamics it is thus essential to disentangle how cells respond to their environment from how they affect it. However, this cannot easily be done using traditional batch culture techniques.

Here, we use a novel microfluidics-based approach that allows us to disentangle how changes in the environment, caused by the collective metabolic activity of a community, affects the growth dynamics of individual cells and how this in turn affects community level activity. Microfluidics allows us to decouple the response of a cell to its environment from the effect it has on the environment itself, something which is not possible to do in conventional experimental approaches. This approach thus allows us to quantify how metabolic interactions between species change over time. Furthermore, we can quantify the effect a species presence has on its own growth dynamics.

We apply our method to study interactions in a naturally derived community that utilizes the polymer chitin. This community consists of a degrader and a cross-feeding species: *Vibrio natriegens* and *Alteromonas macleodii. Vibrio natriegens* is a marine chitin degrader (23). It secretes chitinolytic enzymes into the environment in order to cleave chitin oligomers into monomers and dimers that it can take up into the cell. *Alteromonas macleodii* is unable to degrade chitin (8). Furthermore, it lacks the ability to consume chitin degradation products. As a cross-feeding species, it relies on excreted metabolic by-products produced by the degrader.

## Results and Discussion

### Microfluidics-based approach to study interactions in microbial communities

We developed a novel microfluidics-based approach that allows us to investigate growth dynamics of single cells that experience an environment which is dynamically changing in response to the collective metabolism of a microbial community. In order to achieve this, we couple a mother machine device (24) to a community batch culture and follow cell growth using time-lapse microscopy (Figure 1B). The microfluidic device contains the two members of a simple community consisting of *Vibrio natriegens* - the degrader and *Alteromonas macleodii* - the cross-feeder (Figure 1A). The device is connected to a feeding batch culture in a serum flask. We inoculate bacteria (either the degrader alone or the degrader and the cross-feeder) at low densities into these serum flasks containing sterile growth media and chitin polymer as a carbon source (Figure 1C & D). While microbial cells in the feeding batch culture consume resources and grow, they transition from lag-to exponential to stationary growth phase. In this process they sequentially alter their environment in two major related fashions: the initially available nutrients are consumed by the community members while new nutrients — such as metabolic by-products — become available via the release of metabolites into the environment. The uptake and release of metabolites constantly changes the biochemical environment within the batch culture and consequently in the attached microfluidic device. Cells in the microfluidic device are thus exposed to the same - temporally changing - metabolic environment as the community in the batch culture, but as they are located downstream they cannot change the environment. This approach thus allows us to disentangle how metabolic interactions within microbial communities change over time. Previous studies have used similar approaches in order to answer fundamental questions about microbial physiology in clonal populations (25) (26) (27) (28). Here, we use this approach to study the growth dynamics of individual cells in simple microbial communities.

**Figure 1:**
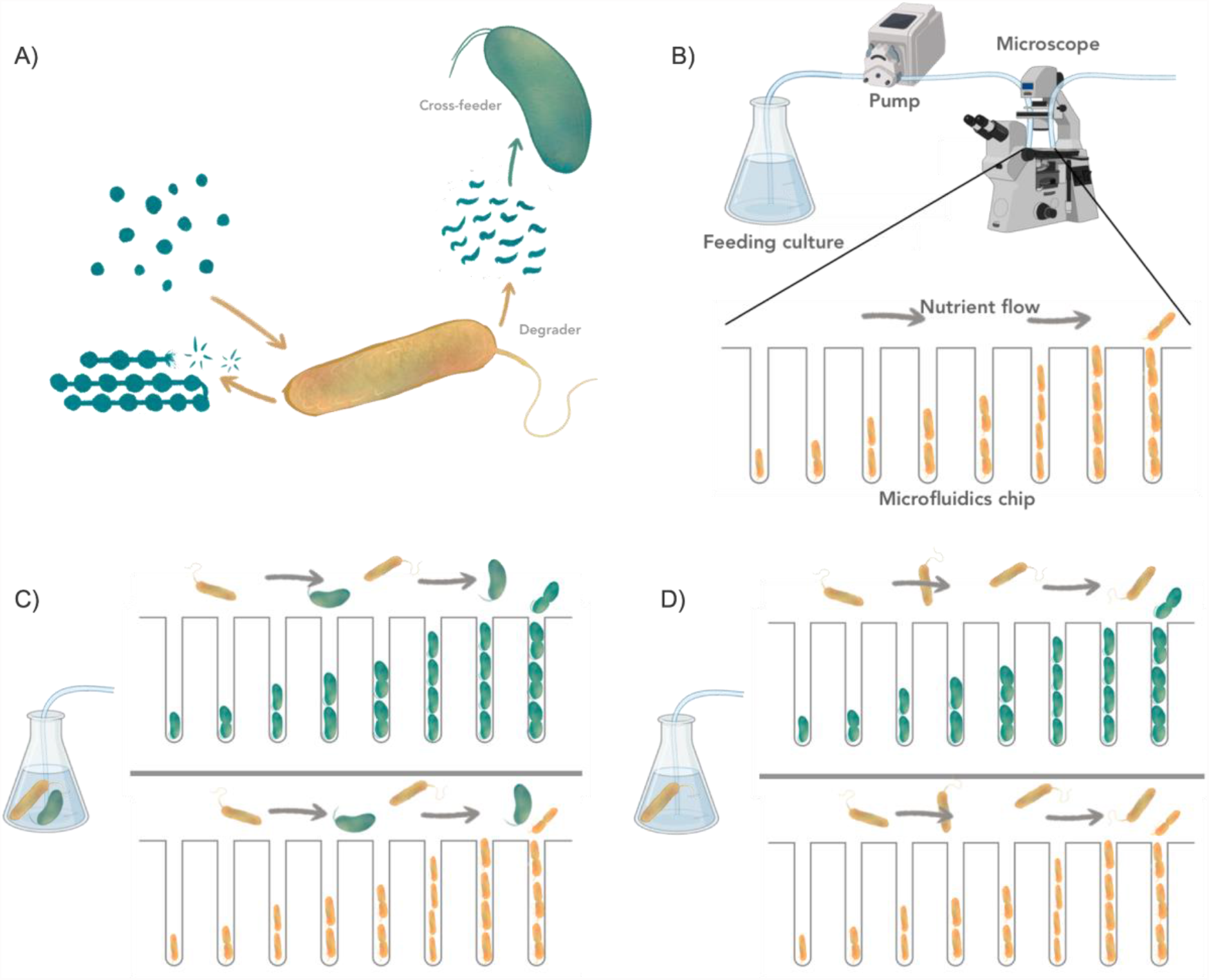
Quantifying the temporal effects of interaction on single cell growth dynamics. (A) Schematic illustration of the metabolic interactions between the degrader (V. natriegens, yellow) and the cross-feeder (A. macleodii, green) when they use chitin as their sole carbon source. (B) Microfluidic setup to quantify the growth dynamics of single cells that experience a batch culture environment. Bacteria are grown in a microfluidic device and are monitored using time-lapse microscopy. The device is connected to a batch culture containing nutrients and bacteria. Via a peristaltic pump, a small proportion of the feeding culture is constantly flushed through the microfluidic chip. Single cells in the microfluidic device experience the same environment as the cells in the feeding batch culture. By the consumption of nutrients and excretion of metabolic by-products, cells in the batch culture change the environment. Cells in the microfluidics device will respond to the change in environment, i.e. via changes in growth rate, but cannot change the environment themselves. (C & D) The microfluidic chips consist of several independent main channels. Each channel is loaded with one of the two species that constitute the community and connected to a single batch culture that contains either the full community with both species (C) or only the degrader species (D). By comparing single cell properties from various batch-microfluidics combinations we can study how the presence of species in communities affect others and themselves. Images were adjusted from BioRender and partially provided by Daniel J. Kiviet.

We first studied the growth dynamics in the metabolic environment created by the full cross-feeding community consisting of both the degrader and the consumer. We inoculated the batch culture with both species and followed the growth over a full cycle (i.e. from lag, via exponential, to stationary phase). In the feeding batch culture, the degrader releases lytic enzymes into the environment and consumes the chitin oligomers. The degradation of chitin and the production and consumption of metabolic by-products are discussed below (Figure 4 and 5). During this process the degrader releases metabolic by-products into the environment which are in turn consumed by the cross-feeder. We analyzed the growth dynamics of the individual species by measuring the growth rates of single cells in the microfluidics device. Both the cross-feeder and the degrader start growing nearly instantaneously after the batch culture is inoculated (Figure 2). This indicates that nutrients for cross-feeding are released as soon as the degraders start with the degradation of chitin. As cells grow and nutrients are depleted both species transition into the stationary phase where their growth rates approach zero. For the degrader growth stops around 25h while for the cross-feeder growth continues until 40h (Figure 2). The difference in length of growth phase can be explained by different metabolic niches: while marine polymer degraders tend to be highly specialized for the polymers they cleave, cross-feeders exhibit a wider metabolic niche (29). Being a generalist for metabolic by-products allows the cross-feeders to grow on various metabolites released by degrader during growth on the polymer (30).

**Figure 2:**
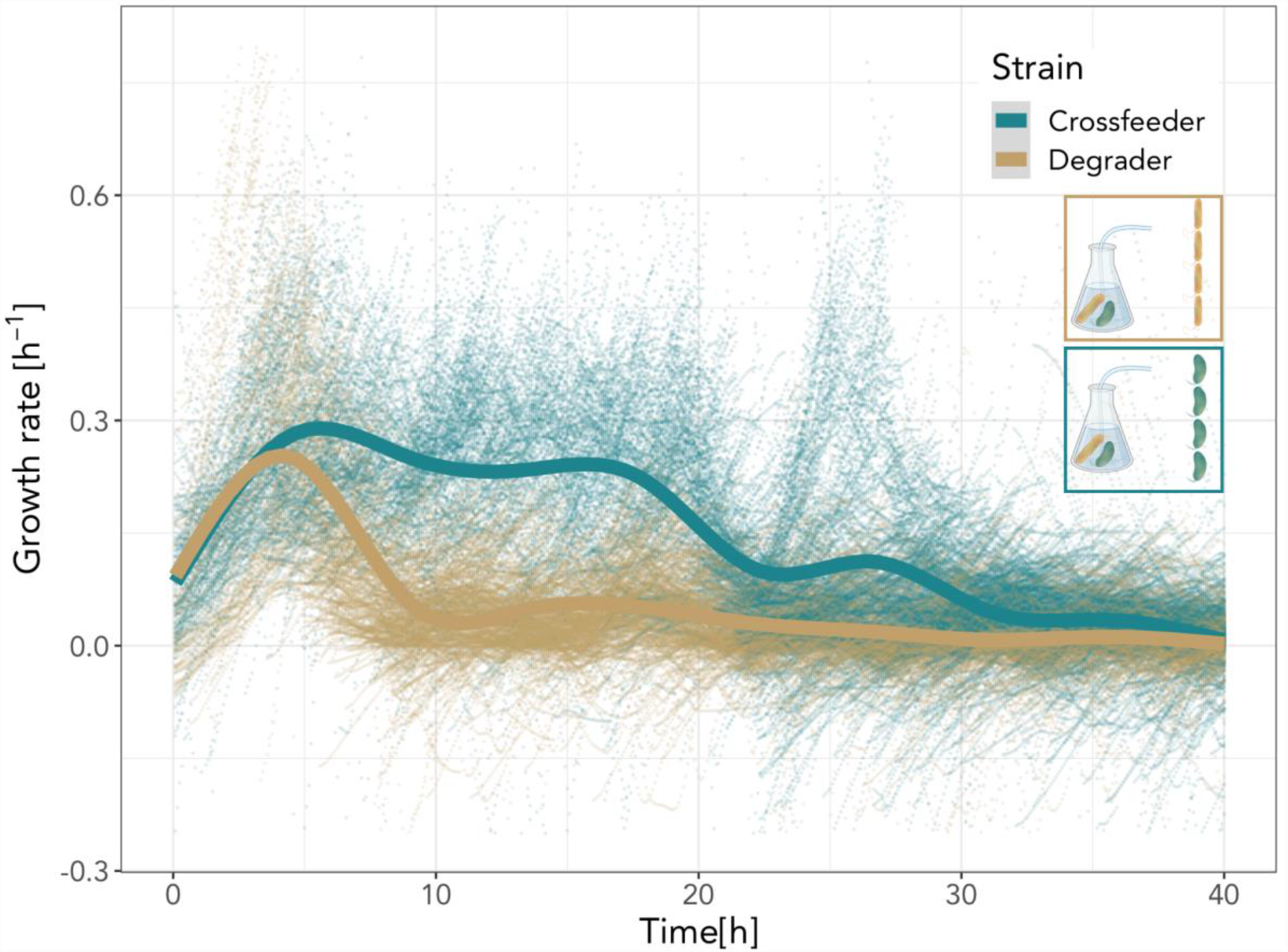
Degrader and cross-feeder cells differ in their growth dynamics when experiencing a co-culture environment. Degrader (yellow) and cross-feeder (green) cells in microfluidic devices were connected to a growing batch culture containing a co-culture of the degrader and the cross-feeder. In the batch culture cells were growing in minimal medium containing 0.1% (w/v) Chitopentaose. Single cell growth rates (points) of each cell present were plotted as a function of time. Lines represent the smoothed means (geom_smooth, ggplot2, RStudio) at each time interval using a generalized additive model. As chitin is degraded via secreted enzymes, resources become available and both cell types start growing. Cross-feeders grow for a longer time before growth rates drop to zero. Four replicate experiments were performed. In total 5608 individual cells were analyzed (N_(Degrader)_ = 1707; N_(Cross-feeder)_ = 3901). This leads to 270’677 instantaneous single cell growth rates (N_(Degrader)_ = 140’126; N_(Cross-feeder)_ =130’551). 1363 (N_(Degrader)_ = 1086; N_(Cross-feeder)_ = 274) data points are not shown because they fall out of the axis range.

### Interactions between degraders and cross-feeders are time dependent

We observed that growth rates changed in a non-monotonic way over time (Figure 2); this raises the question whether metabolic interactions between degraders and cross-feeders change with time and how this affects the growth of the degrader. To address this question, we measured the single cell growth dynamics of degrader cells in two different environments. Specifically, we fed degrader cells with two different batch cultures; one culture contained a co-culture of degrader and cross-feeder—as shown above in Figure 2—while the second culture contained the degrader in mono-culture.

We found that the growth dynamics of the degrader in these two conditions differed in two major ways (Figure 3A): Initially the growth rate of the degrader was higher in the presence of the cross-feeder. This indicates that the cross-feeder facilitates the degrader’s growth in the initial stages of their metabolic interaction (Figure 3A & 3C). This process also leads to a tendency for a higher overall biomass accumulation for the degrader in the presence of the cross-feeder in the first 20h, although with large variation between replicates (Figure 3B & 3D). At the end of the growth phase however, the overall biomass accumulation tended to be higher in degrader cells that experienced a mono-culture environment without cross-feeders (Figure 3B & 3D). At later stages degrader cells in mono-culture exhibit a clear secondary growth phase (Figure 3A). This growth phase is missing in the co-culture environment. Degrader cells in the mono-culture environment grow at higher rates in this secondary growth phase than individuals that experience a co-culture environment. The secondary growth phase can be explained by the reuptake of previously excreted metabolic by-products and corresponds to the diauxic shift observed in various microbiological systems (31) (32).

**Figure 3:**
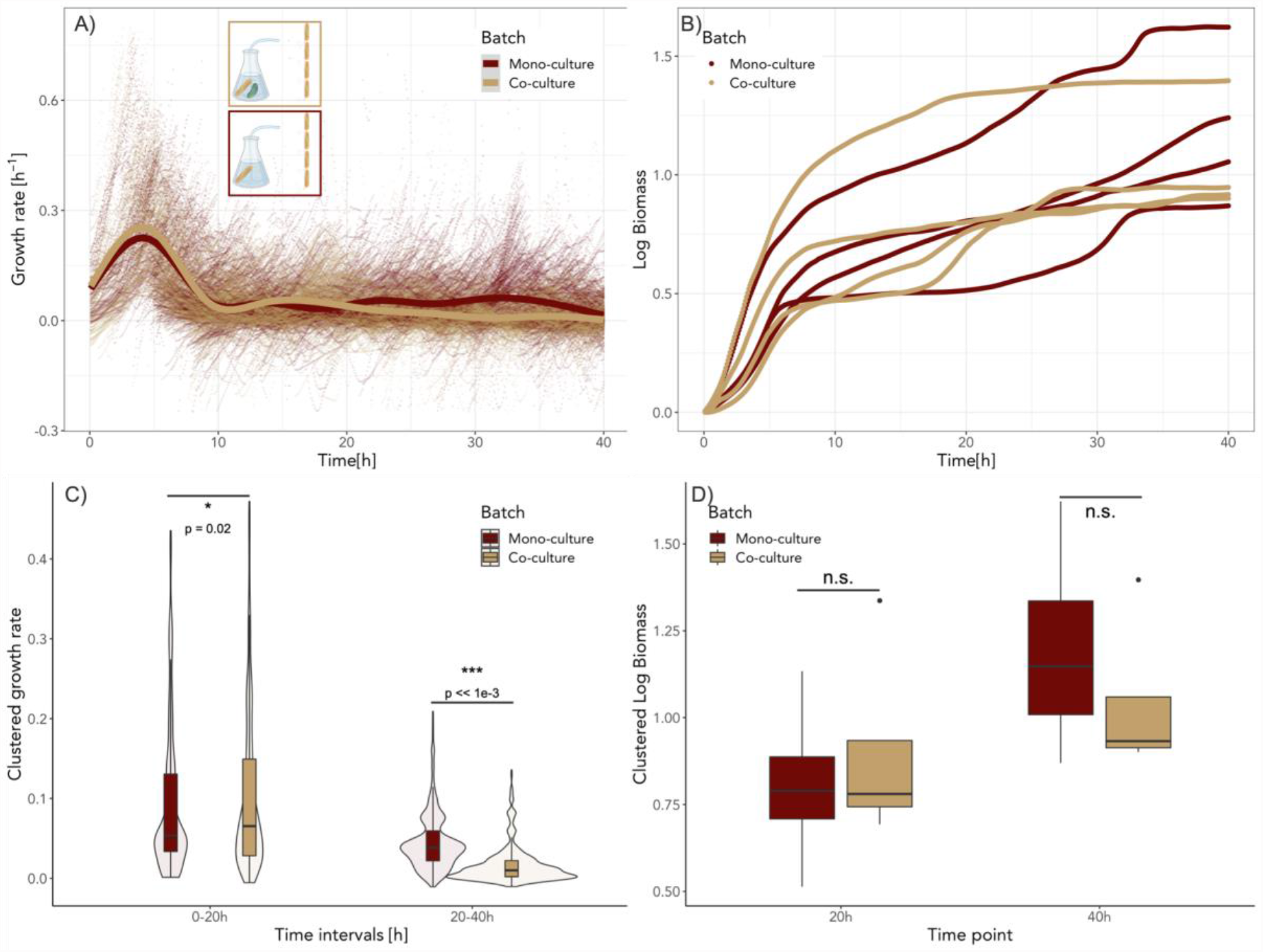
Growth of the degrader is affected by the presence of the cross-feeder in a time dependent manner. Degrader cells in microfluidic chips were connected to two different batch cultures: one with degrader cells in mono-culture (red) and one with a co-culture of degrader and the cross-feeder (yellow). (A) Single cell growth rates (points) are plotted as a function of time. Lines represent the smoothed means (geom_smooth, ggplot2, RStudio) using a generalized additive model. During the first hours the degrader cells supplied with co-culture media grow faster than cells experiencing a mono-culture environment. At later time points we observe a marked secondary growth phase for cells fed by mono-culture that is absent for cells fed by co-culture. Four replicate experiments were performed. Overall 99 mother machine channels were analyzed (N_(Co-culture)_ = 49; N_(Monoculture)_ = 50). In total 3934 individual cells were analyzed (N_(Mono-culture)_ = 2227; N_(Co-culture)_ = 1707). 1494 (N_(Mono-culture)_ = 405; N_(Co-culture)_ = 1089) data points are not shown because they fall out of the axis range. (B) Biomass accumulation of degrader cells when experiencing a mono- or co-culture environment. Cells that experience a co-culture environment (yellow) tend to grow to higher yields in the first 20h than cells that experience a mono-culture environment (red). Due to benefits of the secondary growth phase, cells in mono-culture seem to reach higher overall yields at the end of the experiment. (C) Comparison of average growth rate of degrader cells between the two main growth phases. Single cell growth rates were averaged for each time point (Figure S2) and binned into two 20h time windows. During the first 20h the single cell growth rates were significantly higher when the degrader experienced a co-culture environment (red) compared to a mono-culture condition (Welch Two Sample t-test; mean of co-culture = 0.10, mean of mono-culture 0.09; N_(Co-culture)_ =956; N_(Monoculture)_=956, t = −2.34, p = 0.02). Growth during the secondary growth phase is significantly higher in mono-culture (mean = 0.07 and mean = 0.03 for mono-culture and co-culture respectively, t = 21.15, p < 0.001, N_(Co-culture)_ = 956; N_(Monoculture)_ = 956, Welch Two Sample t-test). (D) Comparison of total cell growth for both batch culture conditions at two time points. For each microfluidic replicate biomass accumulation was calculated (see Methods). During the primary growth phase degrader cells showed no statistical difference in biomass accumulation. Analysis using two-way ANOVA (with the factors culture-type and day of the experiment) revealed no statistically-significant difference in biomass accumulation between co-culture and mono-culture after 20h (f(1) = 2.47, p = 0.21). After 40h analysis using two-way ANOVA (with the factors culture-type and day of the experiment) revealed no statistically-significant difference in biomass accumulation between co-culture and mono-culture (f(1) = 3.35, p = 0.16).

Our data thus reveals a shift in interactions between the degrader and the cross-feeder over ecological time scales: over the course of the growth period the effect of the cross-feeder on the degrader turns from positive to negative. Initially, the cross-feeder facilitates growth, possibly through the removal of growth inhibitory or production of growth promoting metabolites, which positively influences growth of the degrader. At later stages, the presence of the cross-feeder decreases growth of the degrader, possibly because the two types of organisms compete for metabolic by-products. Our data suggests that degrader cells release metabolic by-products into the media during growth on the polymers (first growth phase), an interpretation that we test and support below. Some of these by-products can be consumed by both the degraders and cross-feeders, however the cross-feeder consumes the released by-products constantly, and as a result degrader cells no longer show a second growth phase when cross-feeders are present. These results show that interactions between community members can change over short ecological time scales.

### Interaction is governed by acetate excretion and consumption

These findings raise the question about which metabolite, or which metabolites, are primarily driving the metabolic interaction between the degrader and the cross-feeder. A common interaction mechanism in marine microbial communities is the production and consumption of acetate (33). In order to test whether acetate production and consumption play a role in the interactions between the degrader and the cross-feeder in our system, we measured acetate concentrations of mono- and co-cultures over time. First, we investigated whether acetate levels vary between the mono- and co-culture at different time points. We found that acetate levels change over time in the two batch culture conditions (Figure 4A). In co-culture acetate concentrations reach lower peak levels. This indicates that the cross-feeder is constantly consuming parts of the produced acetate. In the degrader mono-culture acetate accumulates in the early phases and declines at later time points. This indicates that, given the opportunity, the degrader will consume previously released acetate which leads to the secondary growth. Next, we wanted to investigate how acetate levels change according to population size. We found that in early phases of the experiment the acetate concentration relative to the population size (approximated as optical density) is higher than at later time points (Figure 4B). This indicates that acetate production is a metabolic by-product of the initial chitin degradation. Finally, we wanted to verify whether both cell types were able to utilize the accumulated acetate. We found that both members of the community will consume acetate if they are growing on spent media from either mono- or co-culture batch (Figure 4C). Taken together our results indicate that during the initial phases of chitin degradation acetate is produced as a metabolic by-product. In mono-culture the degrader produces acetate in the early phases of the experiment while chitin is still available and consumes acetate when the primary resource has been depleted (Figure S4). In co-culture the cross-feeder constantly consumes part of the acetate. This prevents a secondary growth phase of the degrader in co-culture condition.

**Figure 4:**
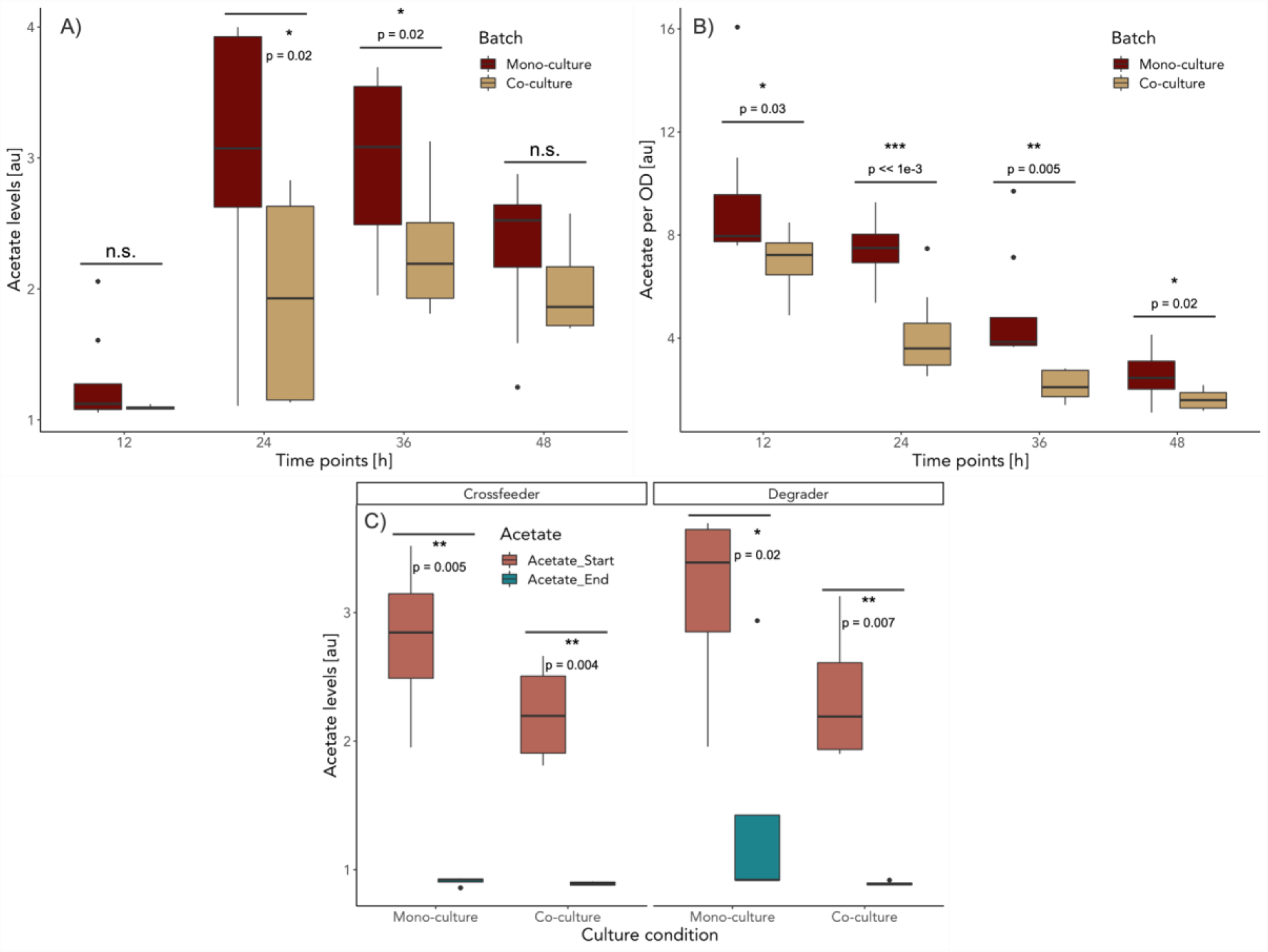
Acetate levels govern cross-feeding interactions. Acetate concentrations measured in batch cultures at various time points. (A) Acetate levels for degrader mono-culture (red) and co-cultures (yellow) in batch cultures change over time. The presence of the cross-feeder leads to lower overall acetate levels. (Welch Two Sample t-test; N = 8; 12h: mean of mono-culture = 1.29, mean of co-culture 1.09; t = 1.50, p-value = 0.09, 24h: mean of mono-culture = 2.95, mean of co-culture 1.93; t = 2.17, p-value = 0.02, 36h: mean of mono-culture = 2.95, mean of co-culture 2.28; t = 2.23, p-value = 0.02, 48h: mean of mono-culture = 2.31, mean of co-culture 2.00; t = 1.27, p-value = 0.11). (B) Acetate levels per OD for degrader mono-culture (red) and co-cultures (yellow) in batch cultures change over time. In the early phases, relative acetate concentration per OD is higher. (Welch Two Sample t-test; N = 8; 12h: mean of mono-culture = 9.39, mean of co-culture 6.94; t = 2.14, p-value = 0.03, 24h: mean of mono-culture = 7.43, mean of co-culture 4.11; t = 4.22, p-value << 1e-3, 36h: mean of mono-culture = 4.96, mean of co-culture 2.15; t = 3.42, p-value = 0.005, 48h: mean of mono-culture = 2.58, mean of co-culture 1.61; t = 2.38, p-value = 0.02). (C) Degrader and cross-feeder cells can consume available acetate. When degrader and cross-feeder are grown on spent media from batch cultures, acetate levels before growth (red) and after 48h of growth (green) vary. (Welch Two Sample t-test; N = 4; Cross-feeder on mono-culture: mean of acetate at start = 2.79, mean of acetate at end 0.91; t = 5.69, p-value = 0.005; Cross-feeder on co-culture: mean of acetate at start = 2.22, mean of acetate at end 0.89; t = 6.50, p-value = 0.004; Degrader on mono-culture: mean of acetate at start = 3.11, mean of acetate at end 2.15; t = 2.61, p-value = 0.02; Degrader on co-culture: mean of acetate at start = 2.35, mean of acetate at end 0.89; t = 5.12, p-value = 0.007).

### Presence of cross-feeder influences degrader’s ability to degrade polymer

In microbial systems that rely on polymer degradation for nutrient availability, growth and enzyme production are tightly coupled: increases in enzyme production and activity increases the concentration of assimilable breakdown products and hence the potential for growth. Since the degraders growth rate is initially increased in the presence of the cross-feeder (Figure 3) we investigated whether the cross-feeders induce enzyme production in the degraders. To this end we measured the enzyme activity when the degrader was growing alone and when it was growing in the presence of the cross-feeder. Using a commercially available enzyme assay kit, we measured the specific activity of chitinases in cell free supernatants. We found that in co-culture chitinase activity is increased when compared to a degrader mono-culture (Figure 5). The cross-feeder does not have any enzymes that cleave the polymer chitin (8); instead our data shows that the presence of the cross-feeder increased the enzyme activity in the degrader cells.

**Figure 5:**
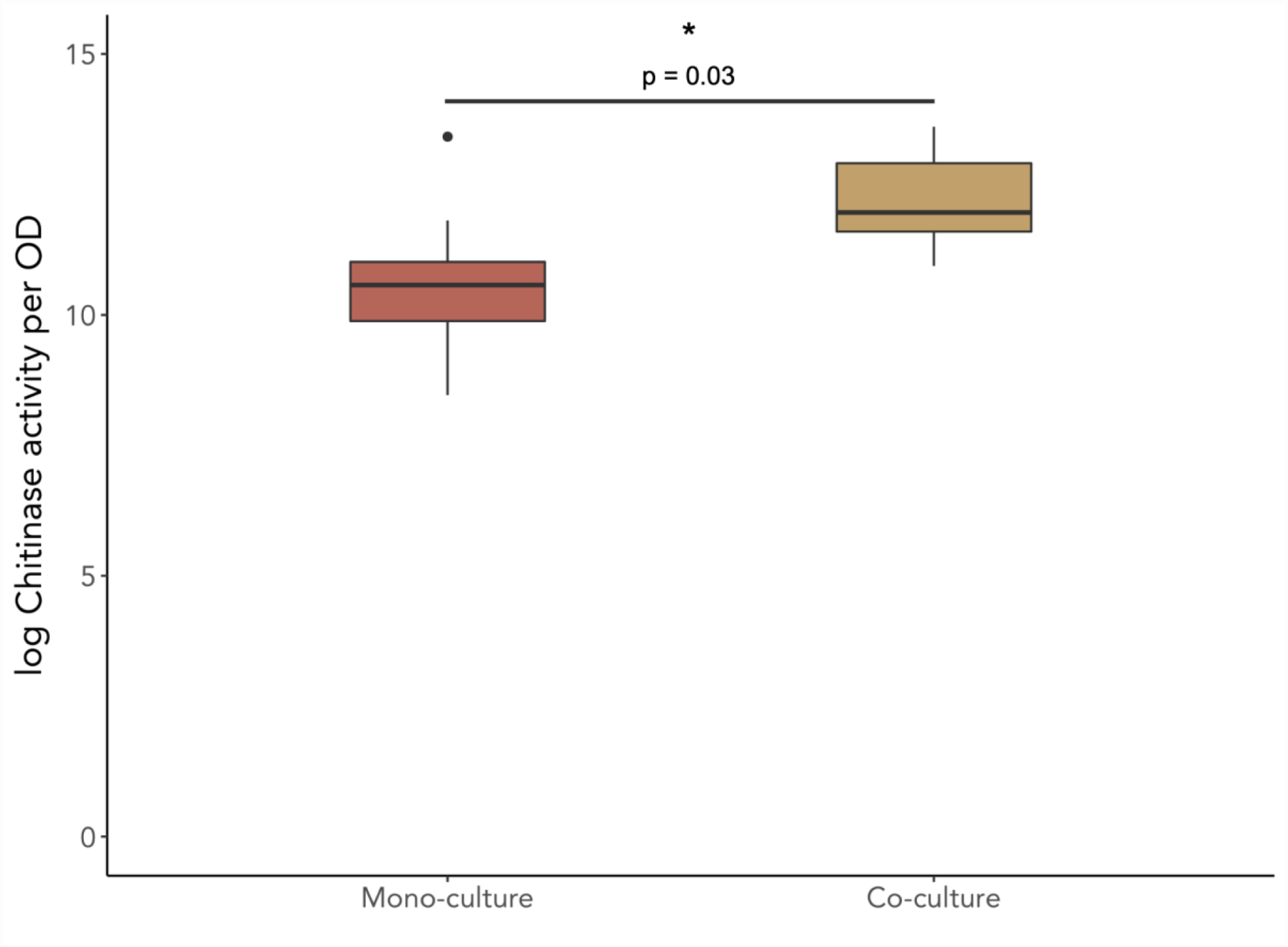
Cross-feeder influences degrader’s chitinase activity. Chitinase activity per unit OD (optical density) in exponential growth phase in minimal media with 0.1% Chitopentaose (w/v). The presence of a cross-feeder significantly increases the overall chitinase activity of the community. (Welch Two Sample t-test on log-transformed data; N = 8; mean of co-culture = 12.21, mean of mono-culture 10.56; t = 2.45, p-value = 0.03).

Higher enzymatic activity can originate in two ways. It is plausible that the effect is a consequence of an increase in enzyme production by the degrader but other effects such as the removal of metabolites (i.e. through cross-feeding) that allosterically hinder the enzymes themselves are also possible. Our data thus suggests that the cross-feeder is able to influence the community level function of chitin degradation by increasing enzyme activity in the degrader. Increased production of lytic enzymes will generally lead to increases in availability of chitin degradation products and therefore higher growth rates for the degrader. The exact mechanism of this phenomenon is still unclear and will be addressed in a future study.

### Effect of cross-feeder on its own growth dynamics

We observed that the cross-feeder increases the growth and chitinase activity of the degrader cells. This raises the question whether the cross-feeders receive an indirect benefit from this interaction, through their promotion of the degrader’s growth and the resulting increase in metabolic by-products. Here we address this question by measuring single cell growth dynamics of cross-feeder cells supplied with nutrients from two different feeding batch cultures. One culture contained a co-culture of degrader and cross-feeder as shown above in Figure 2 while the second culture contained the degrader in mono-culture.

In co-culture the cross-feeder behaves as in a natural community where it can interact with the degrader e.g. by stimulating enzyme production. When connected to a mono-culture, the cross-feeder experiences an environment that is purely shaped by the degrader. Individual cross-feeder cells in the microfluidics device react to metabolic processes of the degrader population in the feeding batch culture without being able to influence them. The natural equivalent of this system would be a community where the cross-feeder is so rare that its presence does not alter the degrader’s behavior while still being able to consume excreted metabolic by-products. Using this approach, we quantify if the cross-feeder grows better as it becomes more numerous and starts to influence the environment.

We find that during the first 20h the positive effect the cross-feeder has on the growth dynamics of the degrader is not reflected in increased biomass production of itself (Figure 6B). When the cross-feeder is part of the community, cells do not solely interact with the degrader but also with conspecifics. This means that in a co-culture environment cross-feeders experience intraspecies competition for the excreted metabolic by-products. This competition is absent in a degrader mono-culture environment. Despite much higher intraspecific competition, biomass accumulation does not decrease in co-culture during the first growth phase (Figure 6D). At later time points, we observe high single cell growth rates and biomass accumulation for the cross-feeder cells that experience a degrader mono-culture condition (Figure 4 A-D). Unlike when they are part of the community, cross-feeder cells grow faster if they are growing on a degrader mono-culture. This difference can be explained by intraspecies competition. Taken together, our data thus suggests that cross-feeders increase the supply of nutrients, but the positive effect is compensated for by the negative effect of increased intraspecies competition.

**Figure 6:**
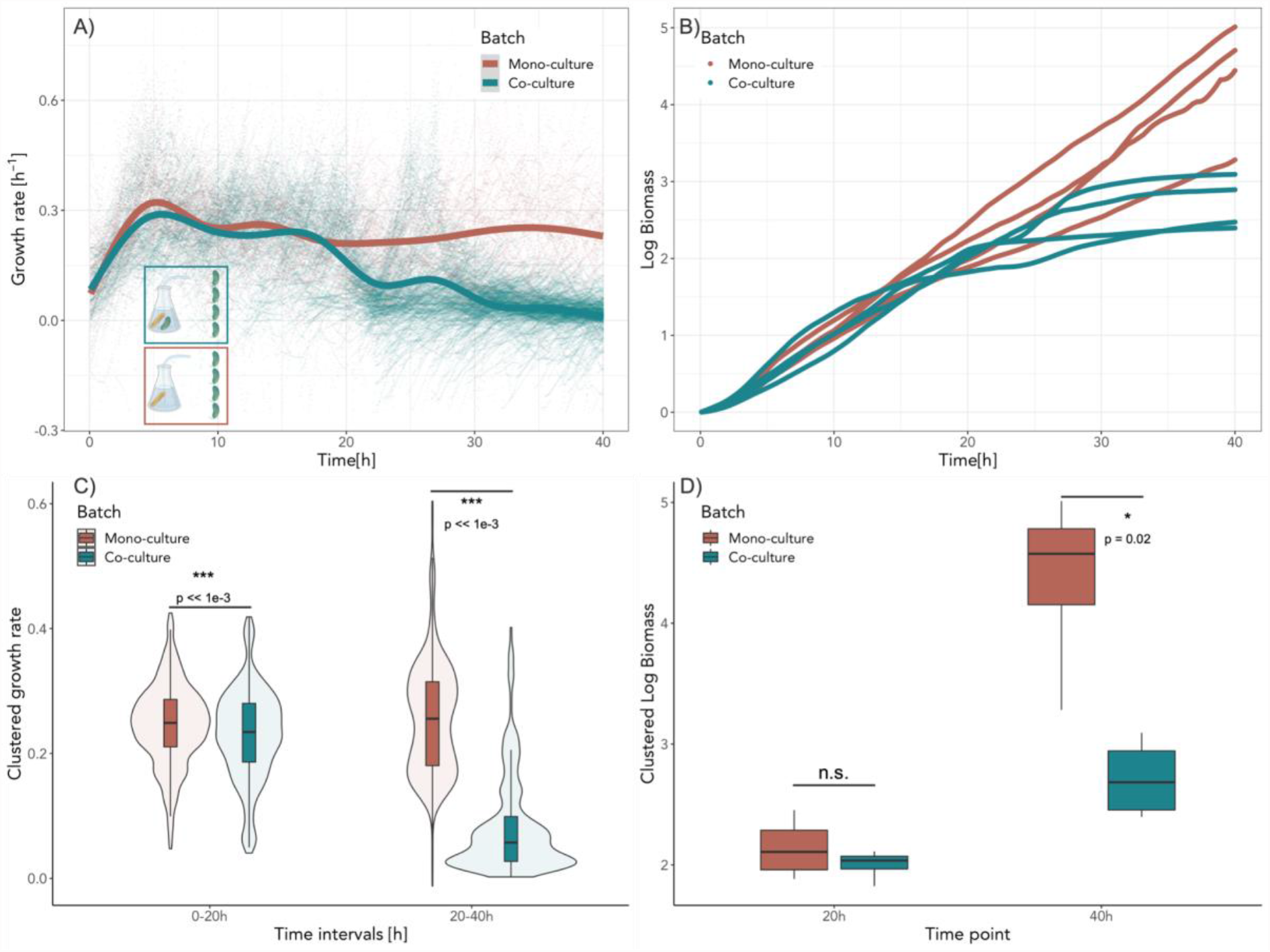
Cross-feeders’ growth dynamics are dominated by intra-species competition. Cross-feeder cells in microfluidic chips were connected to two different batch cultures: one with degrader cells in mono-culture (red) and one with a co-culture of degrader and the cross-feeder (green). (A) Single cell growth rates (points) are plotted as a function of time. Lines represent the smoothed means using a generalized additive model. Initially, cross-feeder cells showed comparable growth trajectories for both environments. As cross-feeder cells in the co-culture consume metabolites excreted by the degrader they use up their available resources and growth rates for cells in the microfluidics device eventual declines. In the degrader mono-culture environment there are no cells that consume all of the previously released metabolic by-products. Under this condition cells in the microfluidics device can grow constantly as they do not experience competition for nutrients. Four replicate experiments were performed. Overall 94 mother machine channels were analyzed (N_(Co-culture)_ = 60; N_(Monoculture)_ = 34). In total 6558 individual cells were analyzed (N_(Mono-culture)_ = 2657; N_(Co-culture)_ = 3901). 322 (N_(Mono-culture)_ = 48; N_(Co-culture)_ = 274) data points are not shown because they fall out of the axis range. (B) Biomass accumulation of cross-feeder cells when experiencing a mono- or co-culture environment. In the first 20h, cells in the two conditions accumulate biomass at the same rate. For cross-feeder cells experiencing a co-culture environment, biomass starts to plateau at the 20h mark. Single cells connected to a degrader mono-culture batch accumulate biomass at a constant rate until the end of the experiment. (C) Comparison of average growth rate of cross-feeder cells between the two main growth phases. Single cell growth rates were binned into two 20h time windows. During the first 20h the single cell growth rates were significantly higher when the cross-feeder experienced a degrader mono-culture environment compared to cells that experienced a co-culture environment. (Welch Two Sample t-test; mean of co-culture = 0.23, mean of mono-culture 0.25; N_(Co-culture)_=956; N_(Monoculture)_=956, t = 4.91, p < 0.001). At later stages growth during the secondary growth phase is significantly higher for cells connected to a degrader mono-culture (Welch Two Sample t-test; mean of mono-culture = 0.26, mean of co-culture 0.08, N_(Co-culture)_=956; N_(Monoculture)_=956, t = 45.31, p < 0.001). D) Comparison of total cell growth for both batch culture conditions for two time intervals. For each microfluidic replicate biomass accumulation was calculated (see Methods). Analysis using two-way ANOVA (with the factors culture-type and day of the experiment) revealed no statistically-significant difference in biomass accumulation between co-culture and mono-culture after 20h (f(1) = 0.61, p = 0.49). During the later stages, cells that experience the mono-culture environment accumulate more biomass. Analysis using two-way ANOVA (with the factors culture-type and day of the experiment) revealed a statistically-significant difference in biomass accumulation between co-culture and mono-culture (f(1) = 21.88, p = 0.02).

**Figure 8:**
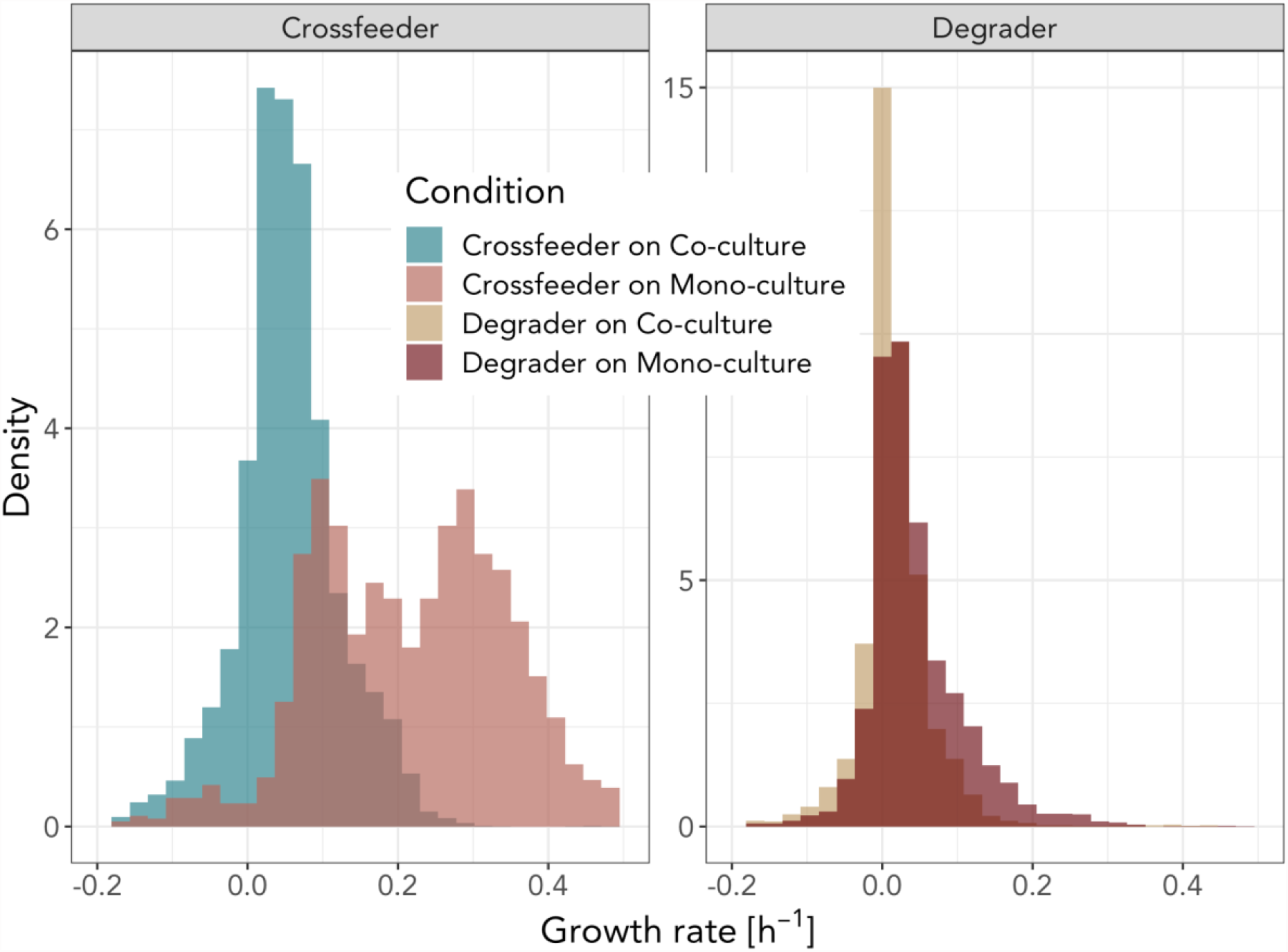
Variation of single cell growth rates. Variation of single cell growth rates differ for degrader and cross-feeder under different conditions. A) Density distributions of single-cell growth rates of cross-feeder on co-culture (green) and mono-culture (light red) between 34h-36h. Coefficient of variation (cv) and variance of distributions (var) show clear differences between these two conditions. Cross-feeder on Mono-culture: cv = 0.61 var = 0.018, Cross-feeder on Co-culture CV = 1.37 var = 0.005. Calculation for bimodality using hartigan’s dip test show a clear bimodal distribution for the cross-feeder when growing on degrader mono-culture (D = 0.02, p < 0.001) but not for growth on co-culture (D = 0.004, p = 0.75). B) Density distributions of single-cell growth rates for the degrader on co-culture (yellow) and mono-culture (dark red) between 3h-36h. Degrader on Mono-culture: cv = 1.50, var = 0.004, Degrader on Co-culture cv = 4.24 var = 0.003. Bimodality calculations reveal no bimodal growth for either condition. (Hartigan’s dip statistics, co-culture: D = 0.003, p = 0.99, mono-culture: 0.003, p = 0.99)

### Distribution of single cell growth rates depend on environment and interactions

Previous studies have shown that cell growth rates can vary strongly between individual cells in a population (34). This variation in growth rate can either be caused by variation in the micro-environments of cells (35), or by stochastic fluctuations (36). One advantage of our microfluidic approach is that we can directly measure this variation in cell growth rates and assess if and how it depends on interspecies interactions.

We investigated whether variation in cell growth rates and the distribution of cell growth rates depends on the interspecies interactions. We found that the degrader has generally narrower distributions than the cross-feeder (Figure 7). Moreover, the shape of the distribution of single cell growth rates of degrader cells did not vary much between growth conditions. In contrast, for the cross-feeder we observed a strong change in the shape of the growth rate distribution between mono- and co-culture conditions at later time points (Figure S9): In mono-culture conditions growth rates vary strongly between cells and shows a clear bi-modal distribution (Figure 7) with one fast and one slow growing subpopulation. This indicates that if the cross-feeder is rare (mimicked by the mono-culture set-up), part of the population can achieve high growth rates late in the growth season; presumably on metabolic by-products that still occur at substantial concentrations. A second part of the population appears to not grow or grows rather slowly. In co-culture conditions this fast-growing subpopulation is not present, and overall cell growth rates follow a more narrow distribution. This emergence of phenotypic heterogeneity in growth rates between genetically identical cells later in the growth cycle is in line with previous observations of how nutrient limitation can promote growth differences in clonal populations (35).

## Conclusion

Our quantitative single-cell measurements of cells in a community context allows us to ask fundamental questions in microbial systems ecology that are usually difficult to experimentally tackle. 1) How is one species’ growth dynamics affected by the presence of another species? 2) How do interactions between species in microbial communities change over time? 3) Do species benefit from the effects they have on others?

We showed that interactions change over ecological times: during early stages cross-feeders increase the growth of degraders—most likely by increasing chitinase activity—while at later stages they decrease the growth of degraders—by outcompeting them for metabolic by-products, such as acetate. Overall, cross-feeders thus had a negative impact on the growth of degraders. Moreover, we showed that the growth of cross-feeders is primarily affected by intraspecies competition for resources; however, in early phases this negative effect can be offset by an indirect positive effect: by increasing the chitinase activity of the degraders.

In natural ecosystems, metabolic interactions are ubiquitous. Frequently, they are observed when cells colonize and degrade natural polymers such as cellulose or chitin where extracellular enzymes are utilized to digest them into subunits that can be taken up into the cell (9). In these environments, non-lytic species cross-feed on metabolic by-products excreted by polymer degraders. Microbial communities that colonize these polymeric particles follow successional dynamics (8). Diverse microbes with varying metabolic capabilities colonize marine particles at different points in time. We studied the cross-feeding that occurs in these natural systems using a small community. We observed that besides long-term successional patterns, short scale interaction changes influence community level properties in these consortia. Our findings pose important questions about the predominant nature of microbial interactions in natural environments. In nature, polymer degrading microbial communities are subjected to chemo-physical factors. Diffusion or local flow in marine ecosystems might prevent the accumulation of excreted metabolic by-products. This would reduce the degrader’s ability to consume previously released metabolites. Therefore, in nature the interactions between degrader and cross-feeder might be more steered towards positive facilitation and syntropy. Furthermore, these observations have important implications on how we study natural interactions in the laboratory: classical microbiological approaches that depend on culturing and plating would have yielded different results on whether the interactions between the degrader and the cross-feeder were positive or negative depending on the time point of probing.

## Material and Methods

### Bacterial strains, media and batch cultures

We used the wildtype strain *Vibrio natriegens* ATCC 14048 and *Alteromonas macleodii* sp. 4B03 isolated from marine particles (8). Strains were cultured in Marine Broth (MB, Difco 2216) and grown overnight at 25°C. 1mL of cell culture was centrifuged (13000rpm for 2min) in a 1.5ml microfuge tube. After discarding the supernatant, the cells were washed with 1ml of MBL minimal medium medium without carbon source. Cells were centrifuged again and the cell pellet was resuspended in 1mL of MBL (marine minimal medium) (37) (33) adjusted to an 0.002 OD600. Cells from these cultures were used for experiments in MBL minimal medium containing 0.1% (weight/volume) Pentaacetyl-Chitopentaose (Megazyme, Ireland). The carbon source was added to the MBL minimum medium and filter sterilized using 0.22μm Surfactant-Free Cellulose Acetate filters (Corning, USA). A total of 500uL of the prepared cultures (250uL + 250uL for co-cultures) were added to 9.5mL of MBL + 0.1% chitopentaose (v/w) in serum flasks. The flasks included a stirrer and were sealed with a rubber seal. Serum flasks were stored on a bench top magnetic stirrer (500rmp) and connected to the microfluidics setup via Hamilton NDL NO HUB needles (ga21/ 135mm/pst 2)

### Microfluidics

Microfluidics experiments were performed as described previously (38) (39) (40). Cell growth was imaged within mother machine channels of 25 × 1.4 × 1.26 μm (length × width × height). Within these channels, cells could experience the batch culture medium that diffused through the main flow channels. The microfluidic device consisted of a PDMS flow cell (50µm/23µm). The PDMS flow cell was fabricated by mixing the SYLGARD® 184 Silicone Elastomer Kit chemicals 10:1 (w/v), pouring the mix on a master waver and hardening it at 80°C for 1h. The solid PDMS flow cell was cut out of the master waver and holes were pierced at both ends of each flow channel prior to binding it to a cover glass (Ø 50mm) by applying the “high” setting for 30 on the PDC-32G Plasma Cleaner by Harrick Plasma. The flow cell was connected via 40mm Adtech PTFE tubing (0.3mm ID x 0.76mm OD) to a Ismatech 10K Pump with 40mm of Ismatech tubing (ID 0.25mm, OD 0.90mm) which again was connected via 80mm Adtech PTFE tubing (0.3mm ID x 0.76mm OD) via a 5mm short Cole-Parmer Tygon microbore tubing (EW-06418-03) (ID 0.762mm OD 2.286mm) connector tubing to a Hamilton NDL NO HUB needle (ga21/ 135mm/pst 2) that was inserted into the feeding culture. During the whole experiment the pump flow was set to 8.35µL/min (0.5mL/h).

### Time-lapse microscopy

Microscopy imaging was done using fully automated Olympus IX81 or IX83 inverted microscope systems (Olympus, Japan), equipped with a 100x NA1.3 oil immersion, phase contrast objective, an ORCA-flash 4.0 v2 sCMOS camera (Hamamatsu, Japan), an automated stage controller (Marzhauser Wetzlar, Germany), shutter, and laser-based autofocus system (Olympus ZDC 1 and 2). Detailed information about the microscopy setup has been described by D’Souza et al 2021 (41). Channels on the same PDMS Chip were imaged in parallel, and phase-contrast images of each position were taken every 5min. The microscopy units and PDMS chip were maintained at room temperature. All experiments were run at a flow rate of 0.1ml h^-1^, which ensures nutrients enter the chamber through diffusion. Four biological replicates were performed. These replicates consist of four independent microfluidics channels (two for each of the strains). These channels were connected to one of two independent batch cultures.

The microscopy dataset consists of 200 mother machine channels; 49 channels for the degrader on co-culture, 51 for the degrader on mono-culture, 40 for the cross-feeder on mono-culture and 60 for the cross-feeder on co-culture.

### Image analysis

Image processing was performed using a modified version of the Vanellus image analysis software (Daan Kiviet, https://github.com/daankiviet/ vanellus), together with Ilastik (42) and custom written Matlab scripts (version 2017_a).

Movies were registered to compensate for stage movement and cropped to the region of growth channels. Subsequently, segmentation was done on the phase contrast images using Ilastik’s supervised pixel classification workflow and cell tracking was done using the Vanellus build-in tracking algorithm.

After visual curation of segmentation and tracking for each mother machine and at every frame growth parameters were calculated using custom written matlab scripts (39)(Dal Co et al 2019). Lengths of individual cells were estimated by finding the cell center line by fitting a third-degree polynomial to the cell mask; then the cell length was calculated as the length of the center line between the automatically detected cell pole positions (see Kiviet et al 2014 for details).

We quantified cell growth by calculating single cell elongation rates r from measured cell length trajectories: L(t)=L(0)·e^(r·t). Cell lengths and growth rates varied drastically over the time course of the experiment; we thus developed a robust procedure that can reliably estimate elongation rates both for large fast-growing cells as well as for small non-growing cells. We first log-transformed cell lengths, which were subsequently smoothed over a moving time window with a length of 5h (60 time points). We used a second order local regression using weighted linear least squares (*rloess* method of Matlab *smooth* function) in order to minimize noise while maintaining sensitivity to changes in elongation rates. Subsequently the instantaneous elongation rate was estimated as the slope of a linear regression over a moving time window of 30min (7 time points). Time points for which the fit quality was bad (*χ*^2^ > 10^−4^) were removed from the analysis (35). All parameters were optimized manually by visually inspecting the fitting procedure of many cell length trajectories randomly selected from across all replicates.

As cells are continuously lost from the mother machine channels it is non-trivial to calculate the total amount of biomass produced in the chip. We thus need to estimate this quantity from the observed single cell elongation-rates. Specifically, we estimated the total amount of biomass produced until a given time point as:

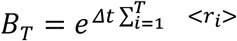

Where < *r*_*i*_ > is the average growth rate of all cells in a given replicate at time point *i*, and where *Δt* is the time interval between two timepoints. By using the average growth rate, we ignore the variation in growth rates between cells; it is, however, highly non-trivial to calculate population growth when growth rates vary both with time and between cells and the current method still allows us to capture the overall effect of interactions on cell growth.

### Datasets and statistical analysis

All microfluidics experiments were replicated 4 times. No cells were excluded from the analysis after visual curation. For *V. natriegens* 2227 cells were analyzed on mono-culture, and 1707 cells were analyzed on co-culture. For *A. macleodii* 2657 cells were analyzed on mono-culture, and 3901 cells were analyzed on co-culture. Each mother machine channel was treated as an independent sample. Statistical analysis was performed in Rstudio v1.2.5033

### Chitinase Assay

Degrader and Cross-feeder cells were cultured in Marine Broth (MB, Difco 2216) and grown overnight at 25°C. 1mL of cell culture was centrifuged (13000 rpm for 2min) in a 1.5ml microfuge tube. After discarding the supernatant the cells were washed with 1ml of MBL minimal medium medium without carbon source. Cells were centrifuged again and the cell pellet was resuspended in 1mL of MBL adjusted to an 0.002 OD600. A total of 10uL of cell culture was added to 190uL of MBL containing 0.1% Chitopentaose (w/v). Cultures were grown to exponential phase in a plate reader (Eon, BioTek) at 25°C. Cell free supernatants were generated by sterile filtering cultures using a multi-well filter plate (AcroPrep) into a fresh 96 well plate. Chitinase activity of cell free supernatants was measured using a commercially available fluorometric chitinase assay kit (Sigma-Aldrich) following the protocol. In short, 10uL of sterile supernatant was added to 90uL of the assay mix. The solution was incubated in the dark at 25°C for 40min before measuring fluorescence (Excitation 360nm, Emission 450nm) in a plate reader (Synergy MX, Biotek). Logarithmic chitinase activity per OD600 was analyzed for 8 replicates.

### Acetate Assay

Cell cultures were prepared and grown in serum flasks as described above. At different time intervals 1mL of culture was removed and OD 600 was measured. Cultures were filter sterilized using 0.22μm Surfactant-Free Cellulose Acetate filters (Corning, USA) into a 1.5mL microfuge tube. Cell free supernatants were stored at −4°C until they were used for acetate measurements. Acetate concentrations were measured using a colorimetric assay kit (MAK086, Sigma-Aldrich) following the protocol. In short, 50uL of cell free supernatant was added to 50uL of assay mix. The solution was incubated in the dark at 25°C for 30min. Acetate concentrations were measured in a plate reader (Eon, Biotek) at 450nm.

### Growth on spend media

Degrader and Cross-feeder cells were cultured in Marine Broth (MB, Difco 2216) and grown overnight at 25°C. 1mL of cell culture was centrifuged (13000 rpm for 2min) in a 1.5ml microfuge tube. After discarding the supernatant, the cells were washed with 1ml of MBL minimal medium medium without carbon source. Cells were centrifuged again and the cell pellet was resuspended in 1mL of MBL adjusted to an 0.002 OD600. A total of 10uL of cell culture was added to the 190uL cell free supernatant described above. Cultures were grown in a plate reader (Eon, BioTek) at 25°C. Cell free supernatants after this growth assay were generated by sterile filtering cultures using a multi-well filter plate (AcroPrep) into a fresh 96 well plate. These supernatants were used as described above to measure acetate levels after growth on spend media.

## Acknowledgments

We thank Ben Roller and Mateus de Oliveira Negreiros for help with the initial data analysis. Alma Dal Co and Susan Schlegel for providing scripts for image analysis; Gabriele Micali, Guga Goram and Glen D’Souza for discussions and advice on analysis of data; Daan Kiviet for designing the microfluidic growth devices; We also thank the members of the Simons Foundation Collaboration on Principles of Microbial Ecosystems collaboration for feedback on the project.

## Funding

MD and MA were supported by the Simons Foundation Collaboration on Principles of Microbial Ecosystems (PriME #542389 and #542395), Swiss National Science Foundation grant 31003A_169978 and by ETH Zurich and Eawag. SvV was supported by Swiss National Science Foundation Ambizione grant PZ00P3_202186.

## Conflict of Interest

The authors declare no conflict of interest.

## Notes

### Competing Interest Statement

The authors have declared no competing interest.

