## Supplementary Figures for "Changes in interactions over ecological time scales influence single cell growth dynamics in a metabolically coupled marine microbial community"

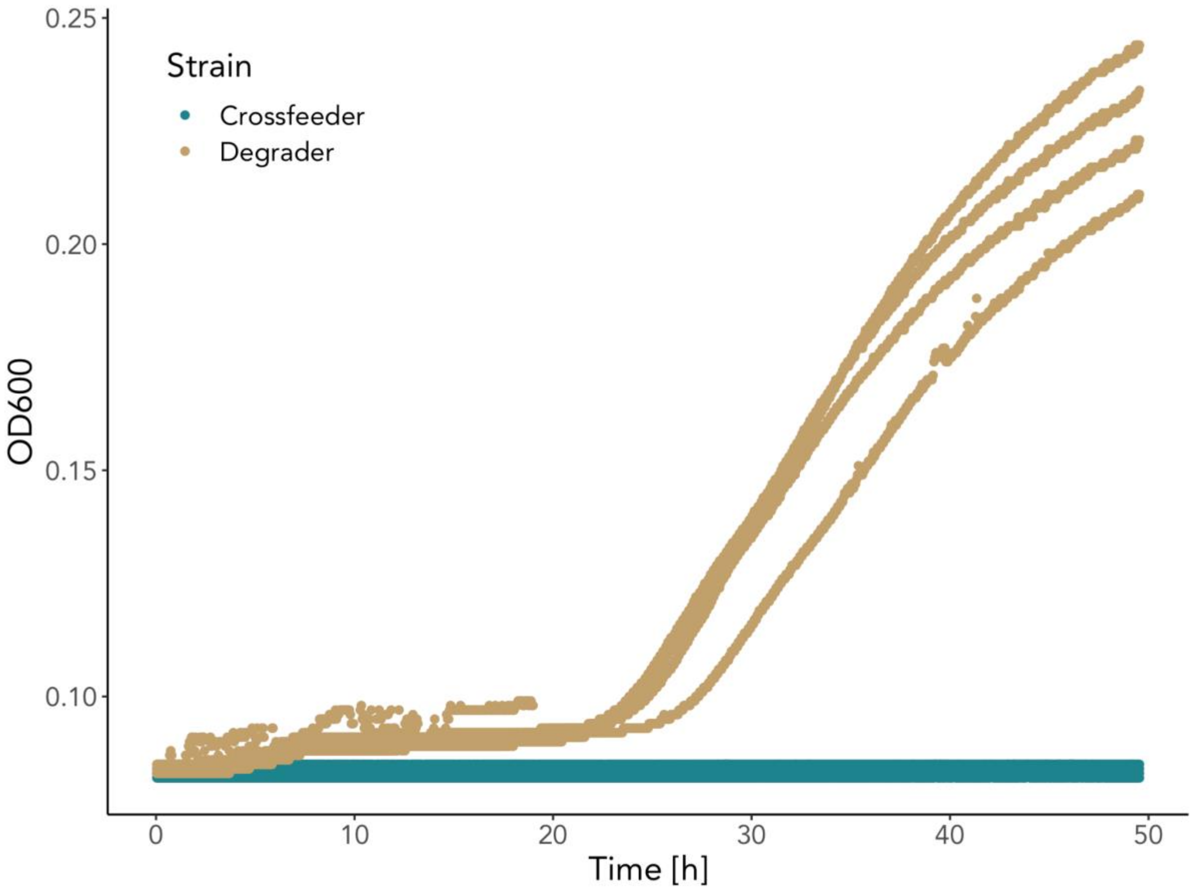


**Figure S1: Growth curve of Degrader and Cross-feeder on Chitin**. Degrader cells release extracellular enzymes and are able to grow on chitin as a sole carbon source (yellow). Cross-feeder cells are unable to grow on chitin (green).

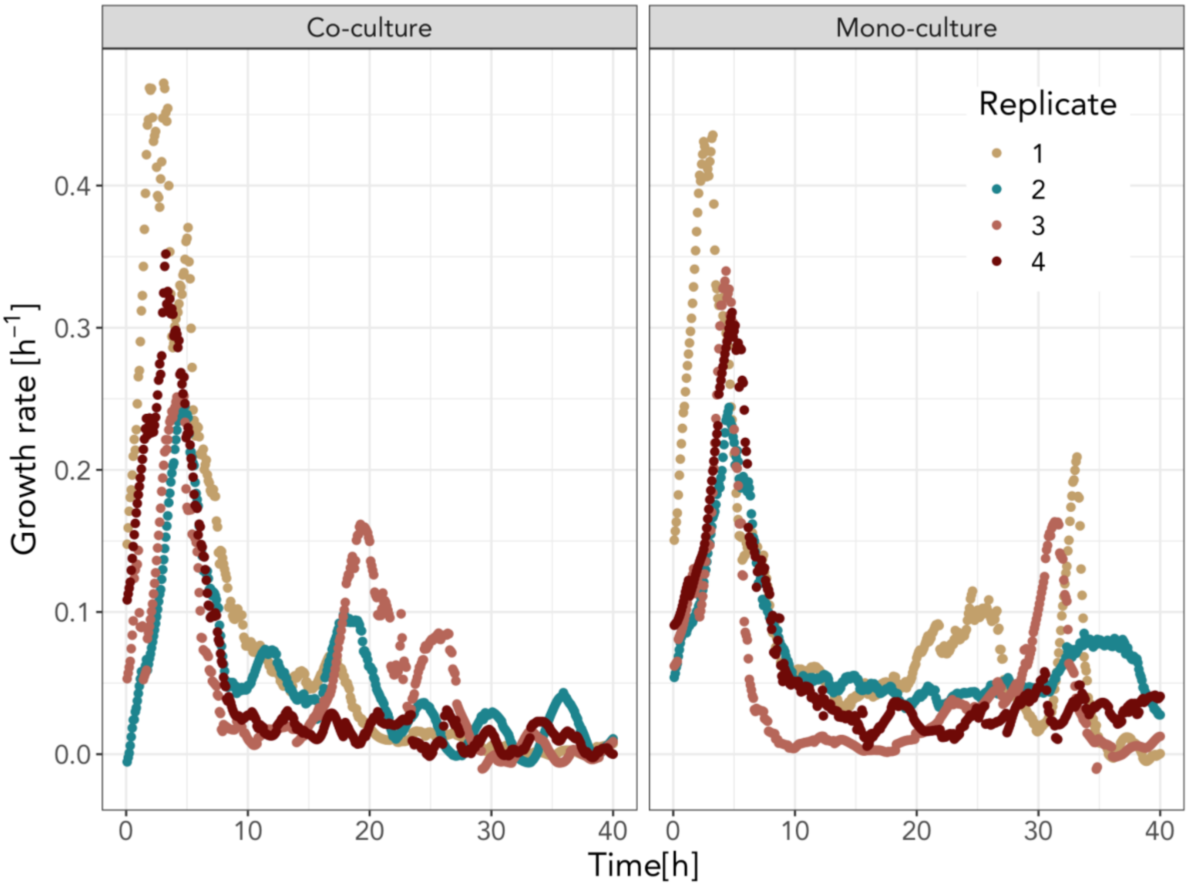


**Figure S2: Mean instantaneous growth rates of degrader cells on mono- and co-culture**. Average growth rates of all single cells at each 5min interval over the 4h growth period of the four replicates. This data was used to calculate differences in average growth rate in the two growth phases (Figure 3).

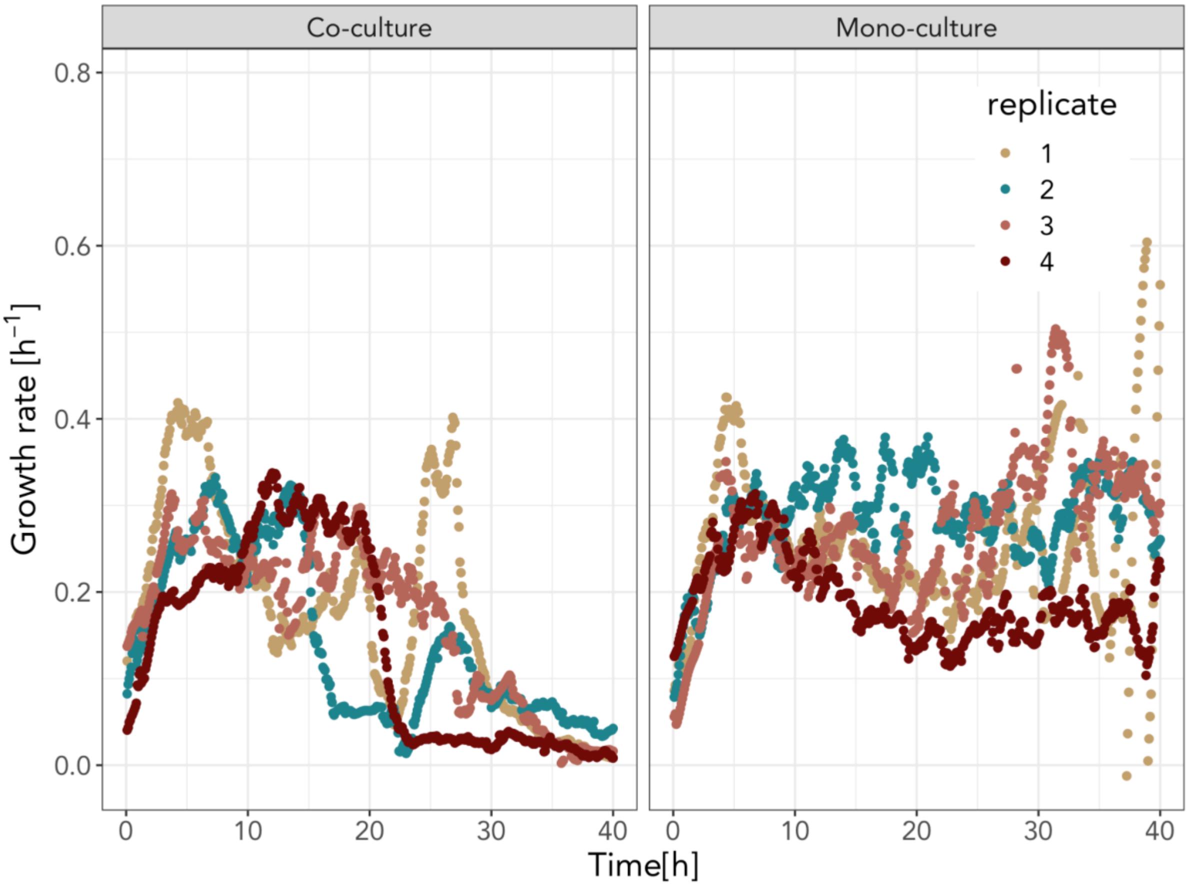


**Figure S3: Mean instantaneous growth rates of cross-feeder cells on mono- and co-culture**. Average growth rates of all single cells at each 5min interval over the 4h growth period of the four replicates. This data was used to calculate differences in average growth rate in the two growth phases (Figure 6).

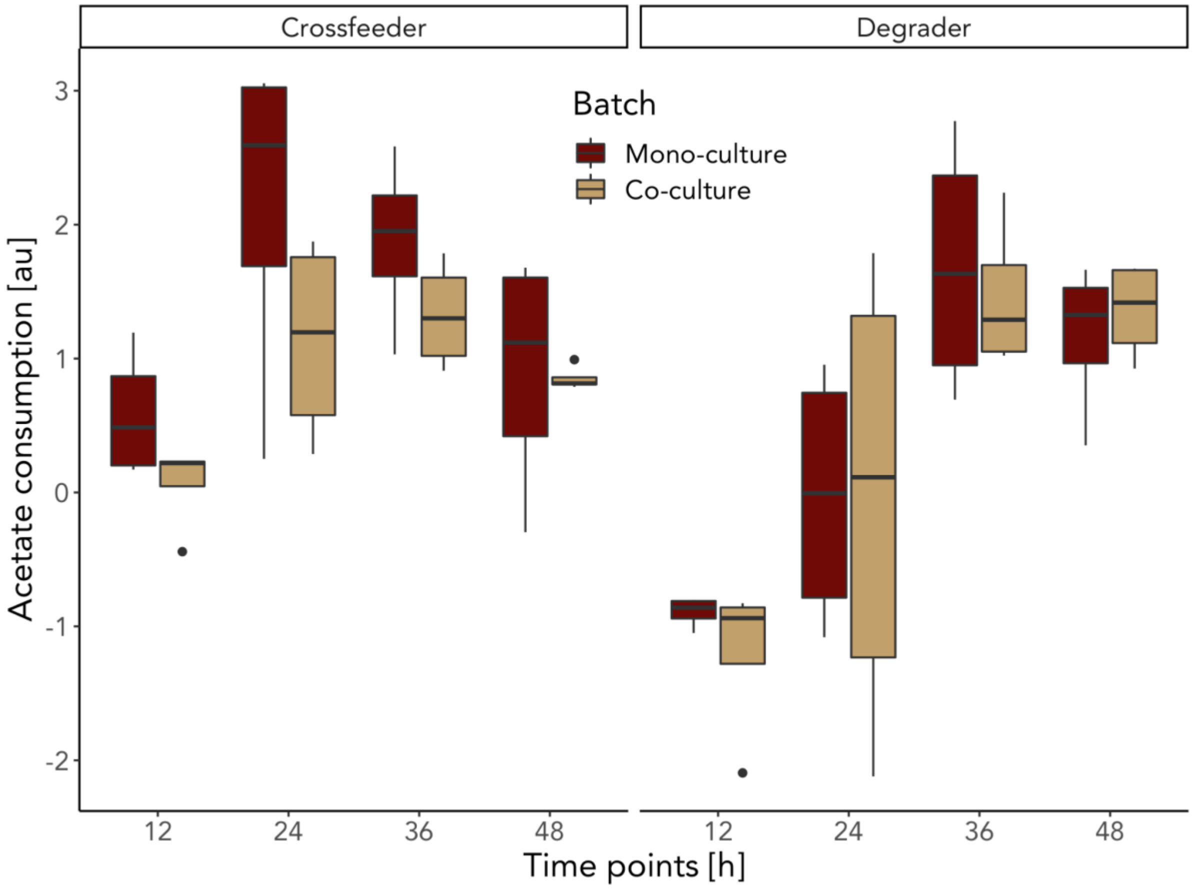


**Figure S4: Acetate consumption at different time points.** Difference of acetate levels before and after the spent media was used as a growth substrate for the degrader or the cross-feeder. Both the degrader and the cross-feeder are able to consume produced acetate (positive values). At early stages when there is still chitin and its primary degradation products in the medium, the degrader will consume the chitin and produce more acetate. This leads to the negative values seen when the degrader is grown on the 12h and 24h spent medium.

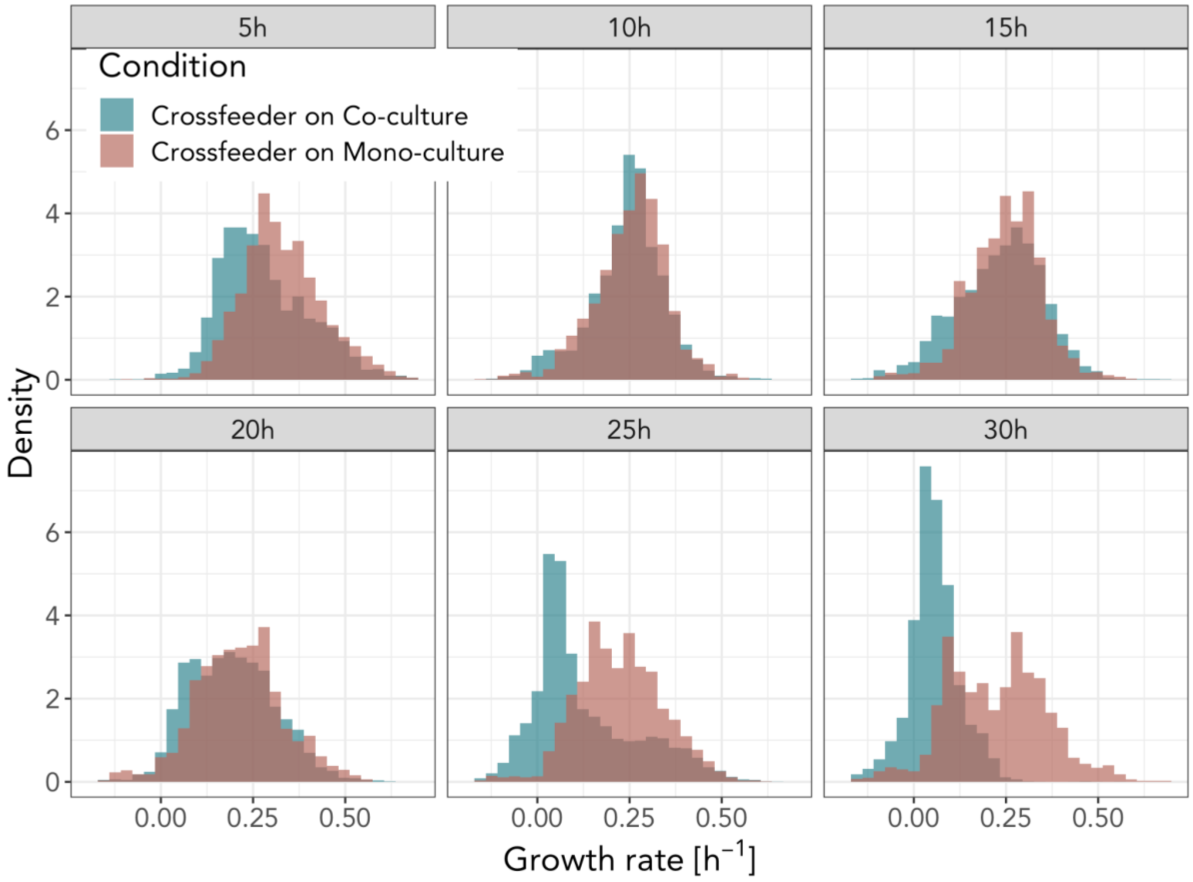


**Figure S5: Single cell distributions for the cross-feeder.** Single cell growth rates of the cross-feeder vary between co-culture (green) and degrader mono-culture (red). At later time points the cross-feeder shows high heterogeneity when experiencing a degrader mono-culture. Distributions are 2h time windows around the indicated time.

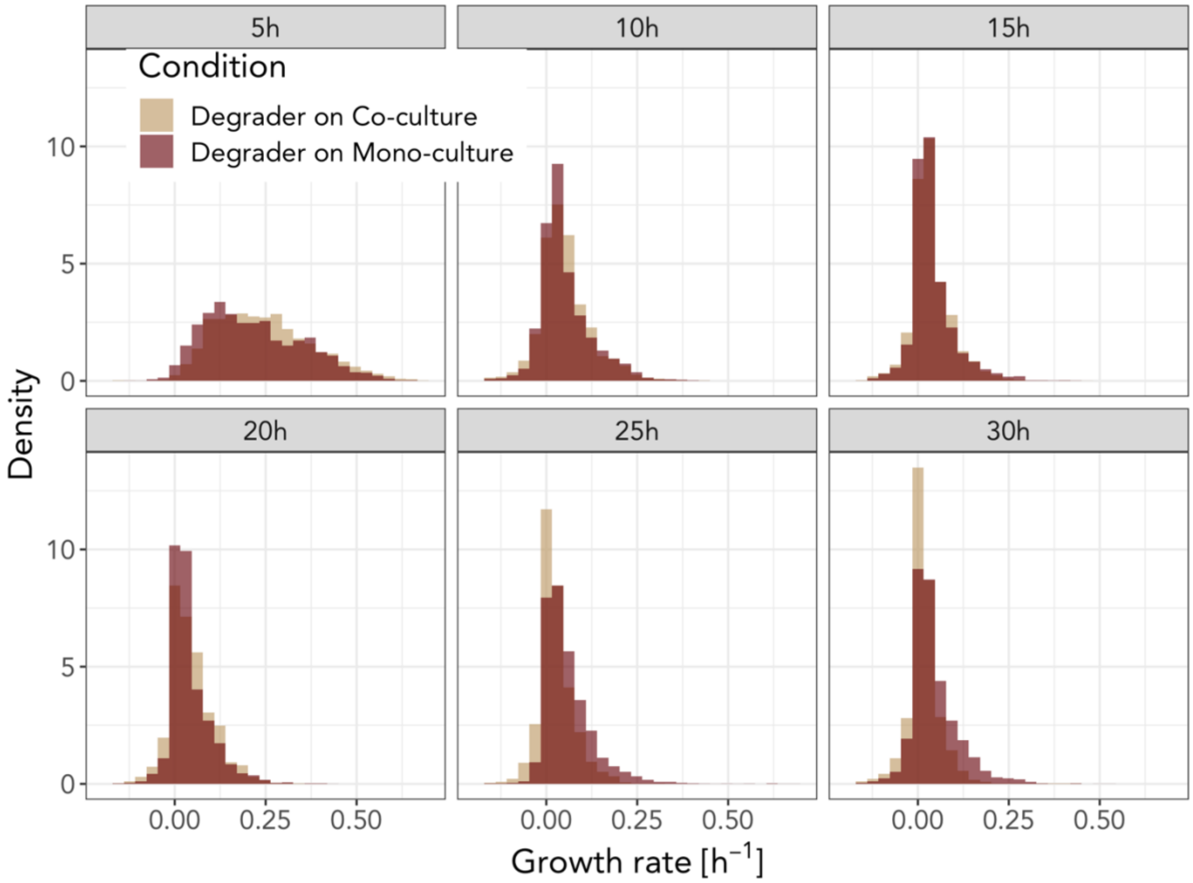


**Figure S6: Single cell distributions for the degrader.** Degrader cells show little heterogeneity in their single cell growth rates during the experiment. Distributions are 2h time windows around the indicated time.

**Single cell growth rates depend on a cell's position in the channel**

We assessed whether variation in micro-environments contributes to variation in cell growth rates. Nutrient availability in the growth channels is determined by the balance of nutrients diffusing into the growth channel from the flow channel and the uptake of these nutrients by cells. When nutrient concentrations in the flow channel are low and uptake rates are high, microscale gradients can be formed. These gradients can affect single cell growth rates across microfluidic channels. In natural environments these gradients can be the result of diffusion from a particulate nutrient source. Cells that are closer to the nutrient source will have immediate access to nutrients while cells that are further away (i.e. in a biofilm surrounding the nutrient core) will be more affected by diffusion and gradients.

In order to investigate which role gradients play in our experiments, we studied the correlation between single cell growth rates and the relative position of individual cells in the channels. We found that for the degrader single cell growth rates were generally higher when cells were closer to the main channel (Figure S5C, S5D). This effect is constant over the period of the experiment (Figure S6). The cross-feeder's growth rate is not affected by the position in the microfluidic device. This indicates that degrader and cross-feeder are generally differentially affected by growth in the microfluidic device, suggesting that the nutrients that the degrader needs are likely available at low concentrations.

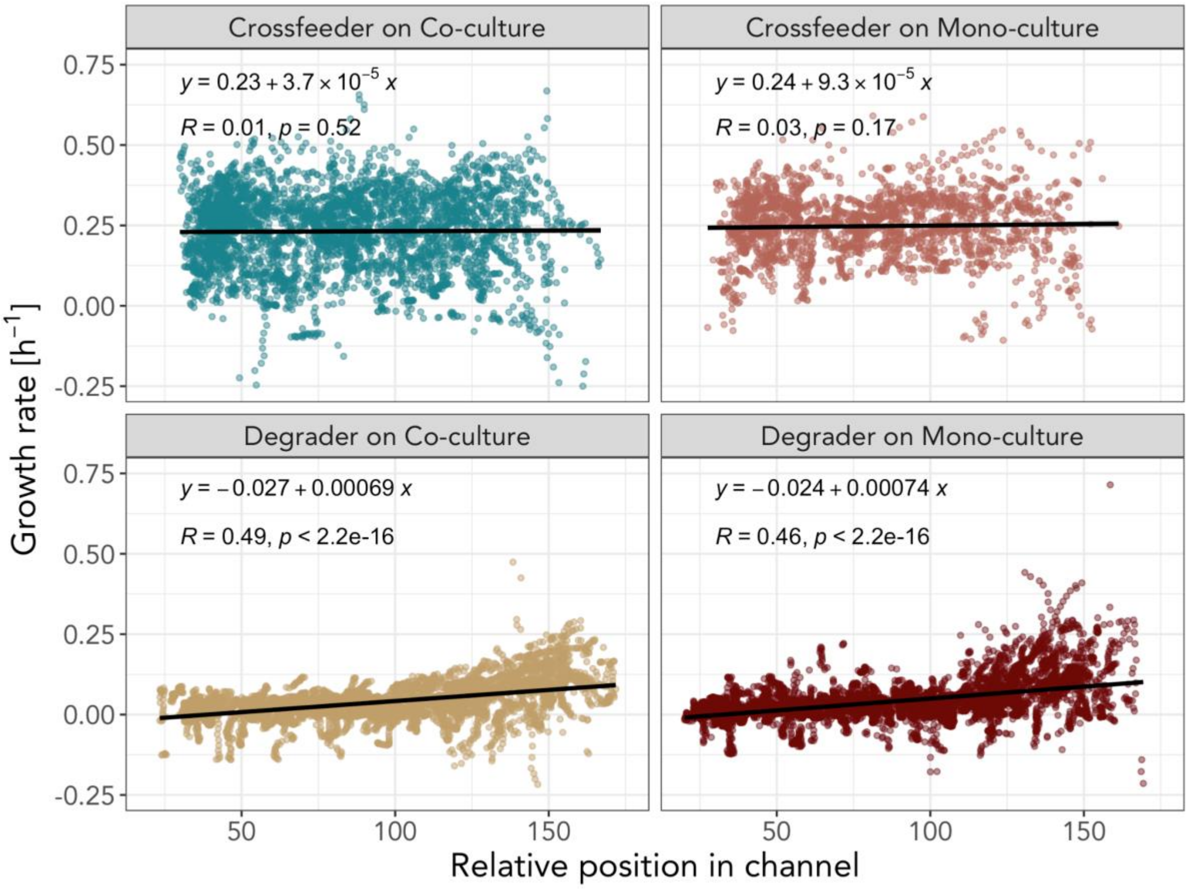


**Figure S7: Position in the growth channel influences single cell growth rates of the degrader.** Degrader and cross-feeder cells are differently influenced by their position within the growth channels. Single cell growth rates between 14h and 16h were plotted against the cell's relative position in the channel. Higher X-axis numbers (relative position in the channel) indicate cells being closer to the main channel. (A) The cross-feeder when grown on co-culture (green) shows no positional effects ($R = 0.01$, $p=0.52$). (B) The cross-feeder when grown on degrader mono-culture (light red) shows no positional effects ($R = 0.03$, $p=0.17$). (C) The degrader when grown on co-culture (yellow) shows a correlation between single cell growth rate and a cell's position in the microfluidic device ($R = 0.49$, $p < 0.001$). (D) The degrader when grown on mono-culture (dark red) shows a correlation between single cell growth rate and a cell's position in the microfluidic device ($R = 0.46$, $p < 0.001$).

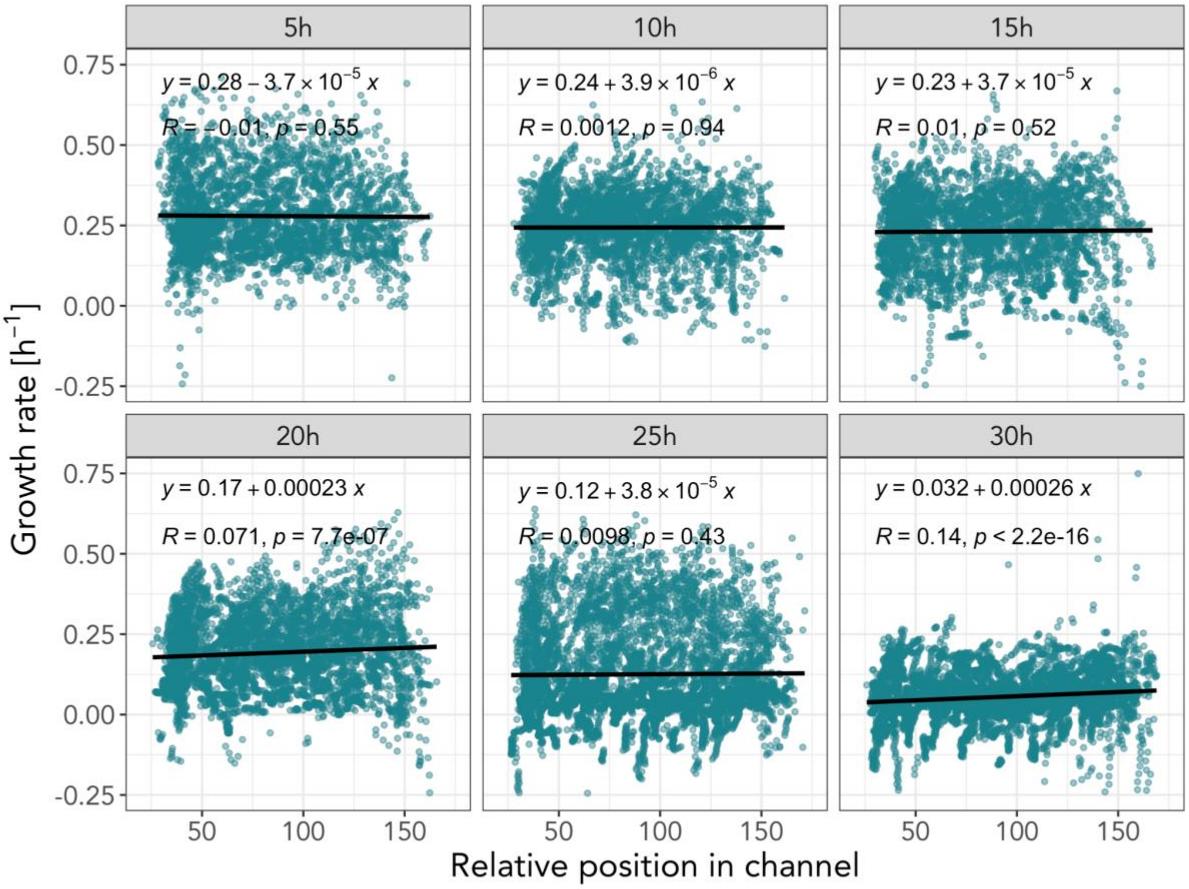


**Figure S8: Single cell growth rates show little variation with position in the growth channel.** Cross-feeder cells that experience a co-culture show almost no correlation in single cell growth rate according to their position in the channel. Single cell growth rates were binned in 2h time windows around the indicated time.

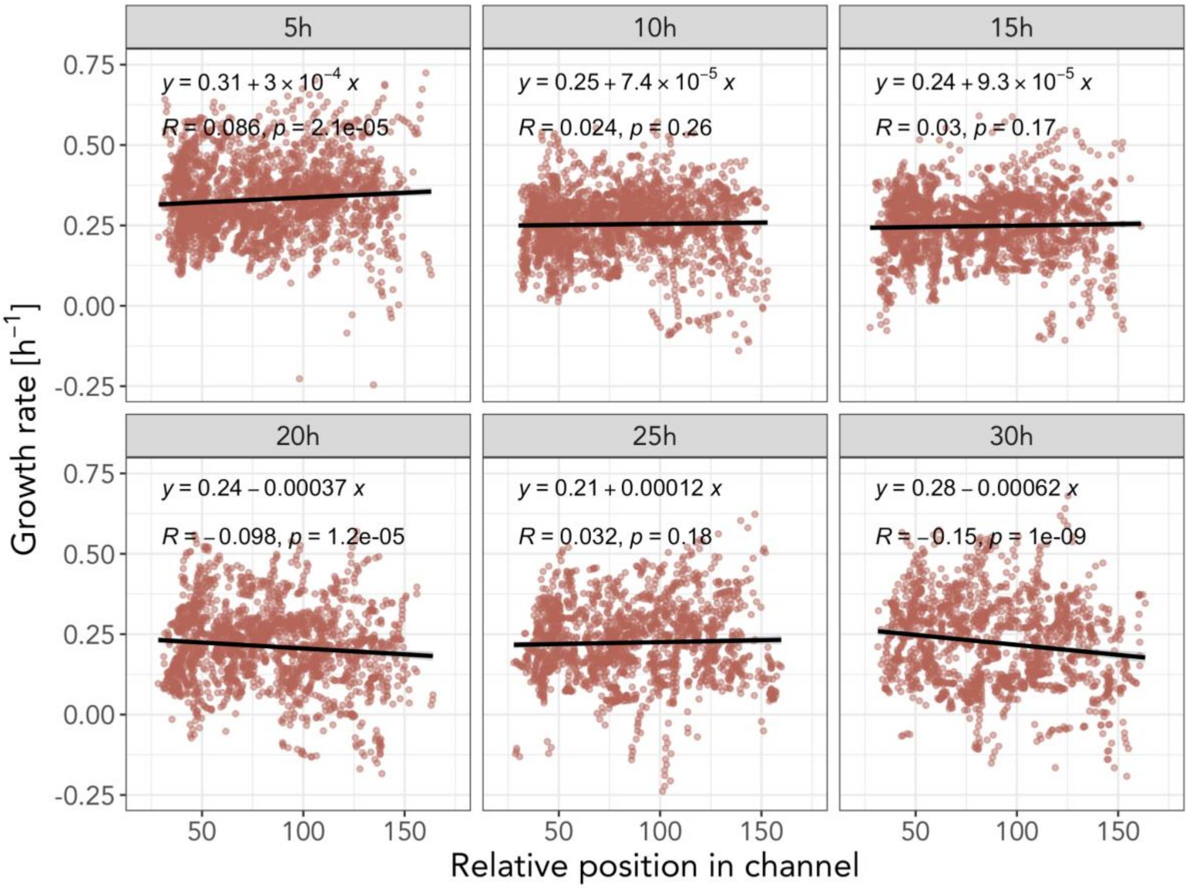


**Figure S9: Single cell growth rates show little variation with position in the growth channel.** Cross-feeder cells that experience a degrader mono-culture show little correlation in single cell growth rate according to their position in the channel. Single cell growth rates were binned in 2h time windows around the indicated time.

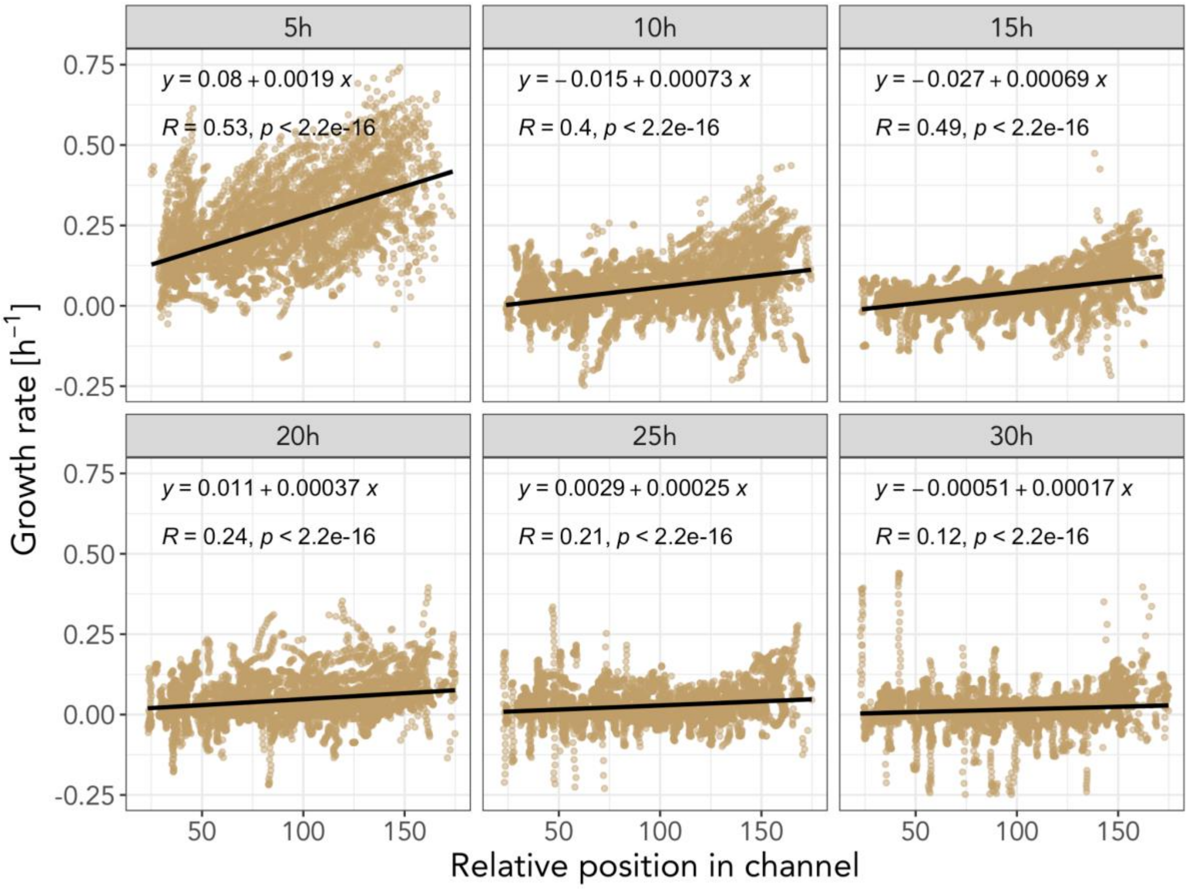


**Figure S10: Single cell growth rates are influenced by the position of cells in the microfluidic channel.** Degrader cells that experience a co-culture show moderate correlation in single cell growth rate according to their position in the channel. Cells closer to the main channel tend to grow generally faster. This effect is consistent over time. Single cell growth rates were binned in 2h time windows around the indicated time.

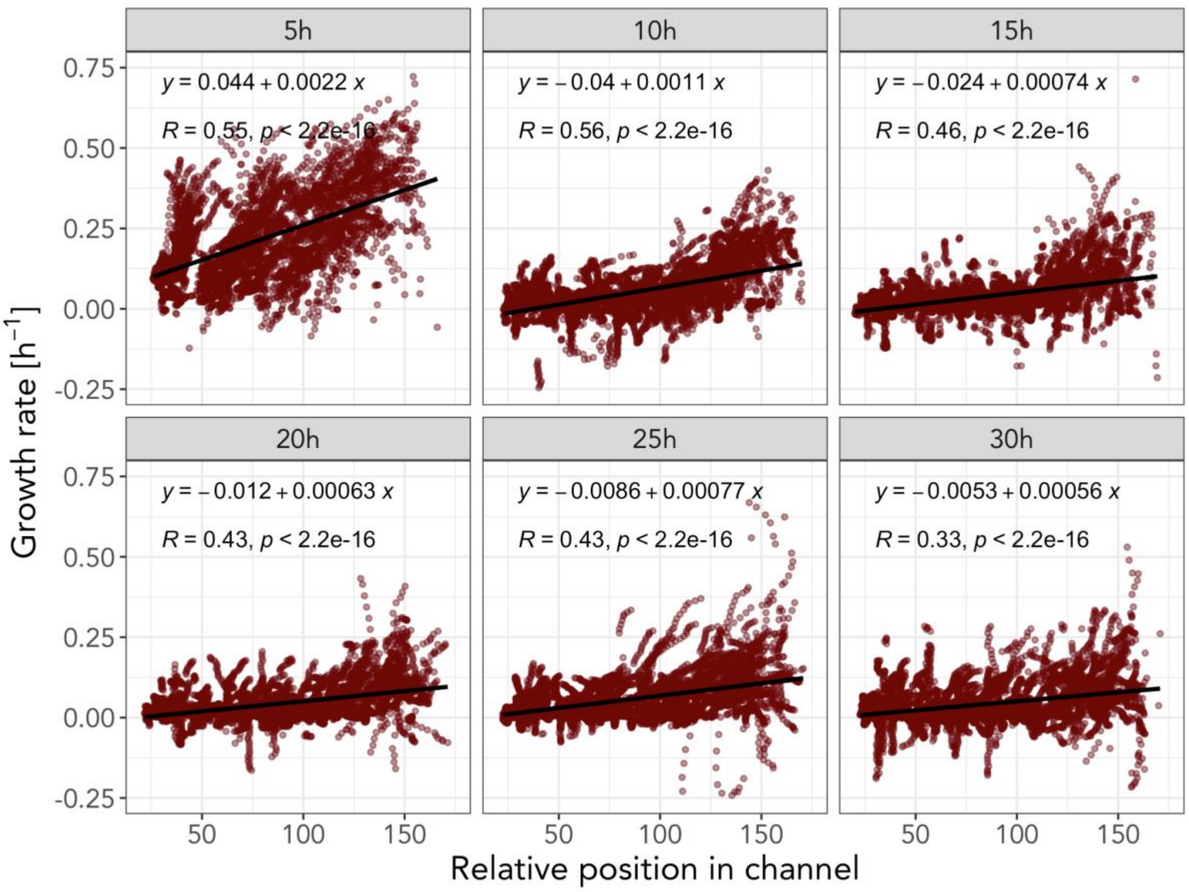


**Figure S11: Single cell growth rates are influenced by the position of cells in the microfluidic channel.** Degrader cells that experience a mono-culture show high correlation in single cell growth rate according to their position in the channel. Cells closer to the main channel tend to grow generally faster. This effect is consistent over time. Single cell growth rates were binned in 2h time windows around the indicated time.
